# The G_2_ phase controls binary division of *Toxoplasma gondii*

**DOI:** 10.1101/2023.07.31.551351

**Authors:** Lauren M. Hawkins, Chengqi Wang, Dale Chaput, Mrinalini Batra, Clem Marsilia, Danya Awshah, Elena S. Suvorova

## Abstract

Division of apicomplexan parasites differs drastically from the division of their host cells. A fraction of apicomplexans divides in the traditional binary mode, such as *Toxoplasma gondii* in asexual stages, whereas the vast majority instead divide in a multinuclear fashion. Such variety of replication modes and a dearth of conserved conventional regulators have hindered the progress of apicomplexan cell cycle studies. We previously identified five Cdk-related kinases (Crk) involved in endodyogenic division of *T. gondii* tachyzoites. The current study investigates the roles of a novel essential cell cycle kinase TgCrk4. We identified this kinase cyclin partner and demonstrated that TgCrk4 regulates processes carried out during conventional G_2_ phase, such as repression of chromosome rereplication and centrosome re-duplication. Accumulation of TgCyc4 in the nucleus and on the centrosomes supported the role of TgCrk4-TgCyc4 complex as a coordinator of chromosome and centrosome cycles in *T. gondii*. Examination of the TgCrk4-deficient tachyzoites confirmed a cell cycle stop prior to the TgCrk6-regulated spindle assembly checkpoint. Furthermore, we identified an ortholog of the DNA replication licensing factor Cdt1 that was a dominant interactor of the TgCrk4-TgCyc4 complex. *T. gondii* Cdt1 is highly divergent but preserved critical signature domains and appeared to play a minimal or no role in licensing DNA replication in G_1_ phase. Functional analyses indicated the primary role of TgCdt1 is in controlling chromosome rereplication and centrosome reduplication. Global phosphoproteome analyses identified immediate TgCrk4 substrates, such as DNA replication licensing factor TgORC4, component of the anaphase-promoting complex TgCdc20, γ-tubulin nucleation factor TgGCP2, and the catalytic subunit of cell cycle phosphatase TgPP2ACA. Importantly, our phylogenetic and structural analyses revealed that the functional TgCrk4-TgCyc4 complex was encoded in the limited group of apicomplexans dividing in a binary fashion. Together with the minimal representation of binary division in Apicomplexa phylum, our findings support the novel view of apicomplexans acquiring binary division to repress ancestral multinuclear mechanisms.

## INTRODUCTION

Apicomplexan parasites are opportunistic intracellular pathogens of humans and animals. They cause many important diseases such as malaria, toxoplasmosis, and cryptosporidiosis. Apicomplexan cell divisions are remarkably versatile and vastly differ from the division mechanisms of their host cells [1, 2]. Some species of apicomplexan parasites duplicate their genome once in a binary fashion, resulting in progenies of two, and others replicate their genomes multiple times, resulting in thousands of daughter cells in a single division round [3]. Duplicated apicomplexan genomes can be segregated after each round of chromosome replication or after multiple rounds, which would include intermediate stages with multiple nuclei, or a DNA syncytium. Depending on the site of daughter bud assembly, parasites can form buds internally or from the surface of the mother cell. There are also species such as *Toxoplasma gondii* that can switch division modes in different hosts [4, 5]. To date, two types of apicomplexan cell division have mainly been studied: the most abundant multinuclear division called schizogony, and the rare binary division endodyogeny.

Cell division is regulated by a cell cycle program, which is poorly understood in apicomplexan parasites. There are several liabilities that prevent fast progress of these studies, such as the complexity of cell division modes, low conservation of cell cycle regulators, and compound parasite-specific internal mitotic structures. Nevertheless, major cell cycle phases and their associated processes have been identified. There are distinguishable growth (G_1_), DNA replication (S), DNA segregation (M or mitosis) phases, and cytokinesis (budding) [2, 6–14]. However, the Gap 2 (G_2_) period that temporarily separates chromosome replication and segregation is presumed missing in apicomplexan cell cycles. To achieve higher progeny, apicomplexan parasites repress cytokinesis and allow chromosomes to re-replicate, which seems to be linked to the binary structure of apicomplexan centrosomes [15, 16]. Mitosis is closed and runs concurrently with the assembly of daughter cytoskeletons (cytokinesis or budding), but multinuclear divisions have additional mitotic mechanisms uncoupled from budding that operate during amplification of nuclear content [8]. Thus, the representative *Plasmodium* spp. engage both mitotic mechanisms to produce multiple daughters, while *T. gondii* tachyzoites involve only coupled mitosis and cytokinesis during endodyogeny [2].

The complete sequence of apicomplexan cell cycle events has never been established and the regulation of cycle transitions remains unknown because only a few major players in the cell cycle pathways of conventional eukaryotes are identifiable in apicomplexan genomes [2, 12]. Most *T. gondii* CDK-related kinases (Crks) and cyclins have limited similarity to conventional eukaryotic counterparts, but we have recently discovered that multiple Crks are required to progress through the *T. gondii* tachyzoite cell cycle [6]. Mapping *T. gondii* Crk activities revealed several points of regulation, including conventional spindle assembly checkpoint (SAC) regulated by the novel TgCrk6-TgCyc1 complex [11]. While G_1_ kinase TgCrk2, S-phase kinase TgCrk5, and mitotic kinase TgCrk6 are conserved among apicomplexans, not all species encode TgCrk4 orthologs [6]. Previous examination of TgCrk4 deficiency using a tet-OFF model suggested a role for TgCrk4 in centrosome duplication, but neither the mechanism nor the cell cycle stage it regulates was determined [6]. We also could not detect a cyclin partner for TgCrk4.

In the current study, we identified the cyclin TgCyc4 that specifically interacts with TgCrk4, and together they regulate the presumably missing G_2_ phase of apicomplexan endodyogeny. We demonstrated that the primary TgCrk4 role is to suppress DNA rereplication and centrosome reduplication. We also found that TgCrk4-TgCyc4 complex interacts with the previously missing ortholog of the. The limited conservation of TgCyc4 
in the genomes of binary dividing parasites, and the specifics of TgCdt1 function suggest the dominant multinuclear division mode was targeted for repression in these apicomplexan subgroups.

## RESULTS

### Coccidian CDK-related kinase TgCrk4 forms complex with atypical TgCyc4

Our analysis of CDK-related kinases in *T. gondii* demonstrated the role of the novel TgCrk4 kinase in dividing tachyzoites [2, 6]. We previously examined TgCrk4 in a tet-OFF model of conditional expression which sufficiently downregulated the kinase after 6-8 hours. Gradual downregulation over a period of one division cycle (7 hours) produced a tainted phenotype and resulted in a rough estimation of TgCrk4-dependent processes. To define processes regulated by TgCrk4, we have now engineered a model of acute proteolytic degradation of TgCrk4. The TgCrk4 auxin-induced degradation (AID) mutant was created by placing an AID-3xHA epitope at the 3’ end of the kinase genomic locus (Fig. S1A and C). Consistent with previous findings, TgCrk4^AID-HA^ kinase was low in abundance and Western blot analysis confirmed downregulation of TgCrk4^AID-HA^ within a 10-min auxin treatment (Fig. 1A). Contrary to the TgCrk4 tet-OFF model, the robust proteolytic degradation of TgCrk4^AID-HA^ resulted in complete growth arrest of the tachyzoites. The RH TgCrk4^AID-HA^ transgenic tachyzoites grew normally in the absence of auxin but failed to form plaques in the presence of auxin (Fig. 1B).

**Figure 1.**
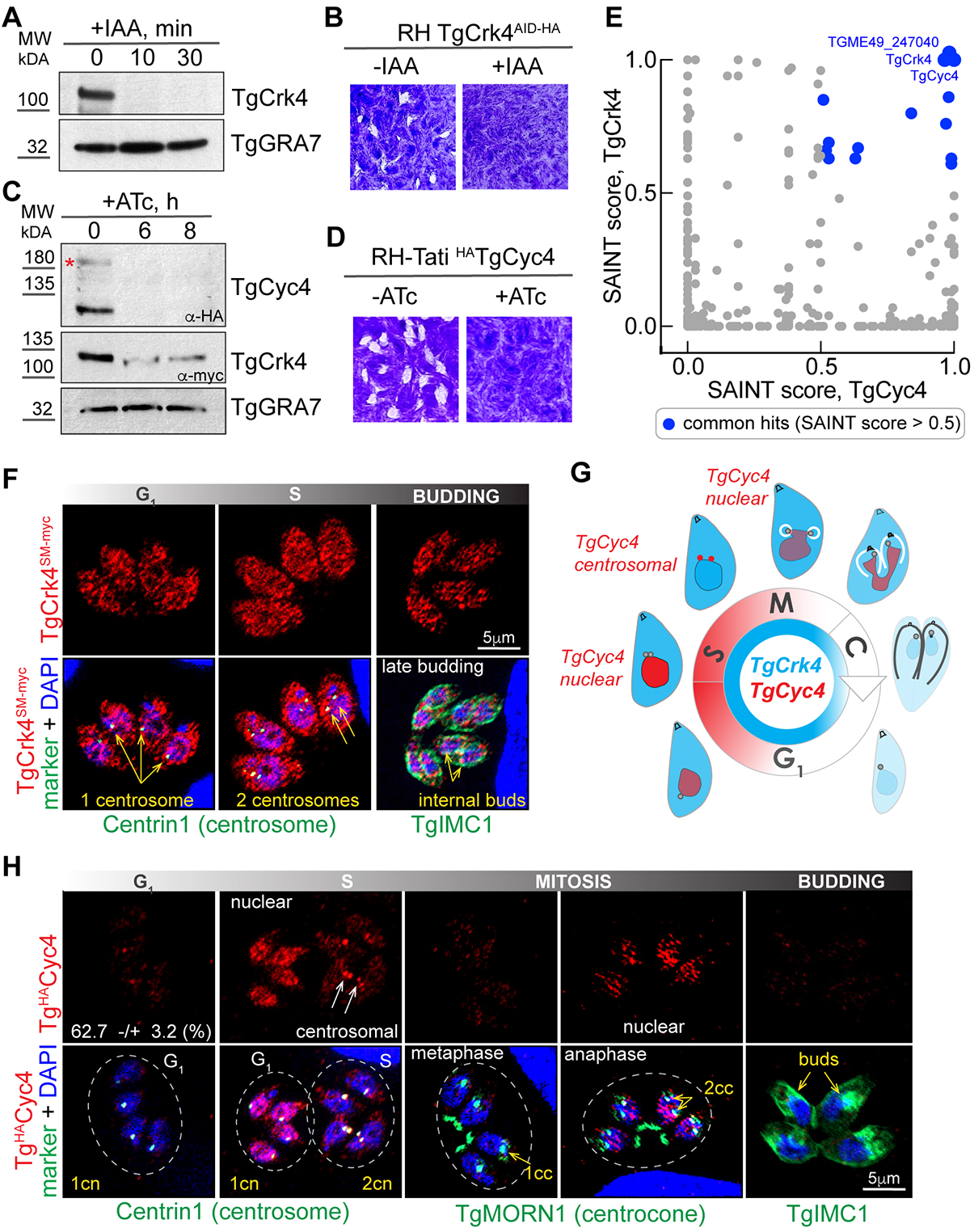
TgCrk4 forms complex with dynamically expressed TgCyc4. (A) The total protein extracts of the RH*ΔKu80TIR1* tachyzoites expressing TgCrk4^AID-HA^ were non-treated or treated with 500μM IAA for 10 and 30 min and analyzed by Western Blot analysis using α-HA (α-rat IgG-HRP) to detect TgCrk4, and with α-GRA7 (α-mouse IgG-HRP) to confirm equal loading of the total lysates. (B) The images of the stained HFF monolayers infected with RH TgCrk4^AID-HA^ tachyzoites and grown with or without 500μM IAA for 7 days. Note that only TgCrk4 expressing tachyzoites (-IAA) formed viable plaques. (C) Western Blot analysis of RH*ΔKu80Tati* tachyzoites co-expressing tetracycline regulatable ^HA^TgCyc4 and TgCrk4^myc^. Total extracts were prepared from parasites grown in the absence or presence of 1μg/ml ATc for 6 and 8 hours. Epitope-fused proteins were detected with α-HA (α-rat IgG-HRP) or α-myc (α-mouse IgG-HRP). The equal protein loading was confirmed with α-GRA7 (α-mouse IgG-HRP) antibodies. (D) The images of the stained HFF monolayers infected with RH*ΔKu80Tati* ^HA^TgCyc4 tachyzoites and grown with or without 1μg/ml ATc for 7 days. (E) Comparison of TgCrk4 and TgCyc4 proteomes using SAINT scores. The common hits with SAINT score >0.5 are highlighted in blue. Note that only three factors, TgCrk4, TgCyc4 and TGME49_247040 had a score of 1, indicating a high probability to form a stable complex. (F) Immunofluorescent microscopy analysis of TgCrk4^SM-myc^ cell cycle expression. The TgCrk4^SM-myc^ (α-myc/α-rabbit IgG Fluor 568) co-staining with Centrin1 (α-Centrin1/α-mouse IgG Fluor 488) was used to distinguish G1 (1 centrosome) and S-phase (2 centrosomes). The budding parasites were identified by co-staining with alveolar protein TgIMC1(α-IMC1/α-mouse IgG Fluor 488). The blue staining represents nucleus (DAPI). Cell cycle phases were determined based on the number of the reference structures and morphology of the nucleus. (G) Schematics of the tachyzoite cell cycle depicts the relative expression of TgCrk4 (blue) and TgCyc4 (red) deduced from immunofluorescent microscopy studies (F and H). Drawings around shows morphological changes of the dividing tachyzoite and associated changes in TgCrk4-TgCyc4 localization. (H) Immunofluorescent microscopy analysis of ^HA^TgCyc4 cell cycle expression. The ^HA^TgCyc4 (α-HA/α-rat IgG Fluor 568) was co-stained with Centrin1 (α-Centrin1/α-mouse IgG Fluor 488) was used to distinguish G_1_ and S-phases and with TgMORN1 (α-MORN1/α-rabbit IgG Flour 488) to identify parasites in metaphase and anaphase of mitosis. The budding parasites were visualized with antibodies against alveolar protein TgIMC1(α-IMC1/α-rabbit IgG Fluor 488). Cell cycle phases were determined based on the number and morphology of the reference structures and morphology of the nucleus (DAPI, blue).

Although TgCrk4 has a recognizable cyclin-binding domain, we previously could not detect a cyclin partner [6]. Phylogenetic analyses of the newly annotated cyclin-domain containing proteins revealed that *Toxoplasma* genomes encoded more cyclin-like factors than initially anticipated [11]. To find out if TgCrk4 interacts with any of these novel cyclin-like proteins, we examined TgCrk4^AID-HA^ protein complexes (Fig. 1E, Fig. S1E). Mass-spectrometry analysis detected a single cyclin-domain containing protein, TgCyc4 (TGME49_249880), and SAINT analysis confirmed a high probability of TgCrk4-TgCyc4 interaction (Fig. S1G, Table S3).

To test whether TgCyc4 is a major partner of the cell cycle kinase TgCrk4, we made several attempts to establish an AID conditional model for TgCyc4. Despite successful incorporation of the AID-3xHA nucleotide sequence at the 3’ end of the *TgCyc4* genomic locus, we could not detect TgCyc4^AID-HA^ protein, suggesting proteolytic processing at the C-terminus. Corroborating this finding, we successfully placed 3xHA epitope at the N-terminus of TgCyc4 and under control of the tetracycline regulatable promoter within the *TgCyc4* genomic locus (Fig. S1B and D). Western blot analysis detected two bands corresponding to the full-length (Fig. 1C, red star) and truncated TgCyc4, further corroborating TgCyc4 processing. This observation, together with the lack of protein expression in TgCyc4^AID-HA^ with a C-terminus tag, suggests that the TgCyc4 C-terminus is highly unstable.

Testing TgCyc4 tet-OFF model, we determined that TgCyc4 was essential for tachyzoites, and a 6-hour treatment with anhydrotetracycline (ATc) efficiently downregulated TgCyc4 (Fig. 1C and D). To confirm TgCrk4-TgCyc4 complex assembly *in vivo*, we isolated ^HA^TgCyc4 complexes and detected TgCrk4 as a major TgCyc4 interactor (Fig. 1E, Fig. S1F and H). To examine whether the complex subunits affect the expression of one another, we introduced TgCrk4^myc^ into TgCyc4 tet-OFF parasites. We found that ATc-induced TgCyc4 downregulation was accompanied by a significant reduction of TgCrk4 expression, suggesting co-dependent stabilization of the TgCrk4-TgCyc4 complex (Fig. 1C). Altogether, our results confirmed that TgCrk4 forms a specific complex with novel cyclin TgCyc4 that is essential for tachyzoite division.

### TgCyc4 expression is dynamic across endodyogeny

We reasoned that if TgCrk4 forms a complex with TgCyc4, then their expression should spatiotemporally overlap. To test this hypothesis, we endogenously tagged TgCrk4 with Spaghetti Monster 10xMyc (SM-Myc) epitope, which significantly improved visualization of this low-abundant kinase. Co-staining with a set of cell cycle markers showed constitutive and ubiquitous TgCrk4 expression with a negligible decline in the tachyzoites’ last (late budding) and first (G_1_) cell cycle stages (Fig. 1F and G, blue schematics). Examining TgCyc4 expression revealed that, contrary to its partner kinase, TgCyc4 was highly dynamic and dramatically changed localization in a cell cycle-dependent manner. Quantification of TgCyc4-positive tachyzoites with single centrosomes (TgCentrin1: single dot in G_1_ cells) showed that TgCyc4 first emerged as a nuclear factor in mid-G_1_ phase (Fig. 1H). This brief period of nuclear expression was followed by a pronounced accumulation of TgCyc4 on duplicated centrosomes (S-phase) that gradually faded by mid-mitosis. Previous studies showed that mitotic marker TgMORN1 facilitates separation of the subphases of mitosis [6, 11]. Besides in the mother basal complex (mBC), TgMORN1 was present on the daughter basal complexes (dBC), known as MORN-rings. In metaphase, dBCs were associated with a single intranuclear compartment centrocone that resolves into two compartments in anaphase (Fig. 1H) [11]. We detected a second peak of TgCyc4 expression at the spindle assembly checkpoint operating during metaphase-to-anaphase transition. Co-staining with TgMORN1 revealed that TgCyc4 briefly accumulated in the nucleus of anaphase cells (2dBCs and two centrocones) and was completely gone by mid-budding (Fig. 1H and G). The spatiotemporal expression of TgCrk4 and TgCyc4 echoed that of conventional cell cycle CDK-cyclin complexes that contain a constitutively expressed kinase and an oscillating cyclin [17]. The alternating cell cycle localization of TgCyc4 suggests that TgCrk4-TgCyc4 complex controls events in the nucleus and on the centrosome.

### TgCrk4 regulates cell cycle events located upstream of the spindle assembly checkpoint

The timing of the TgCrk4-TgCyc4 complex expression indicates its role takes place in post-G_1_ cell cycle phases. To determine the timing of TgCrk4-regulated processes, we compared TgCrk4-dependent cell cycle arrest with the previously examined spindle assembly checkpoint (SAC) regulated by TgCrk6-TgCyc1 complex [11]. Thereby, we established a semi-synchronization approach and monitored the cell cycle progression of parasites enriched at a specific checkpoint. First, we determined the minimal period of TgCrk4 knockdown that parasites tolerated without losing viability. We found that, like TgCrk6, about 80% of parasites reverted to division after 4 hours of TgCrk4 deficiency (+IAA), while a longer treatment severely affected parasite survival (Fig. S2A). Western blot analysis verified the reversibility of this auxin-induced block; it showed that TgCrk4-and TgCrk6-deficient parasites could restore their expression within 1 hour of auxin removal (Fig. 2A).

**Figure 2.**
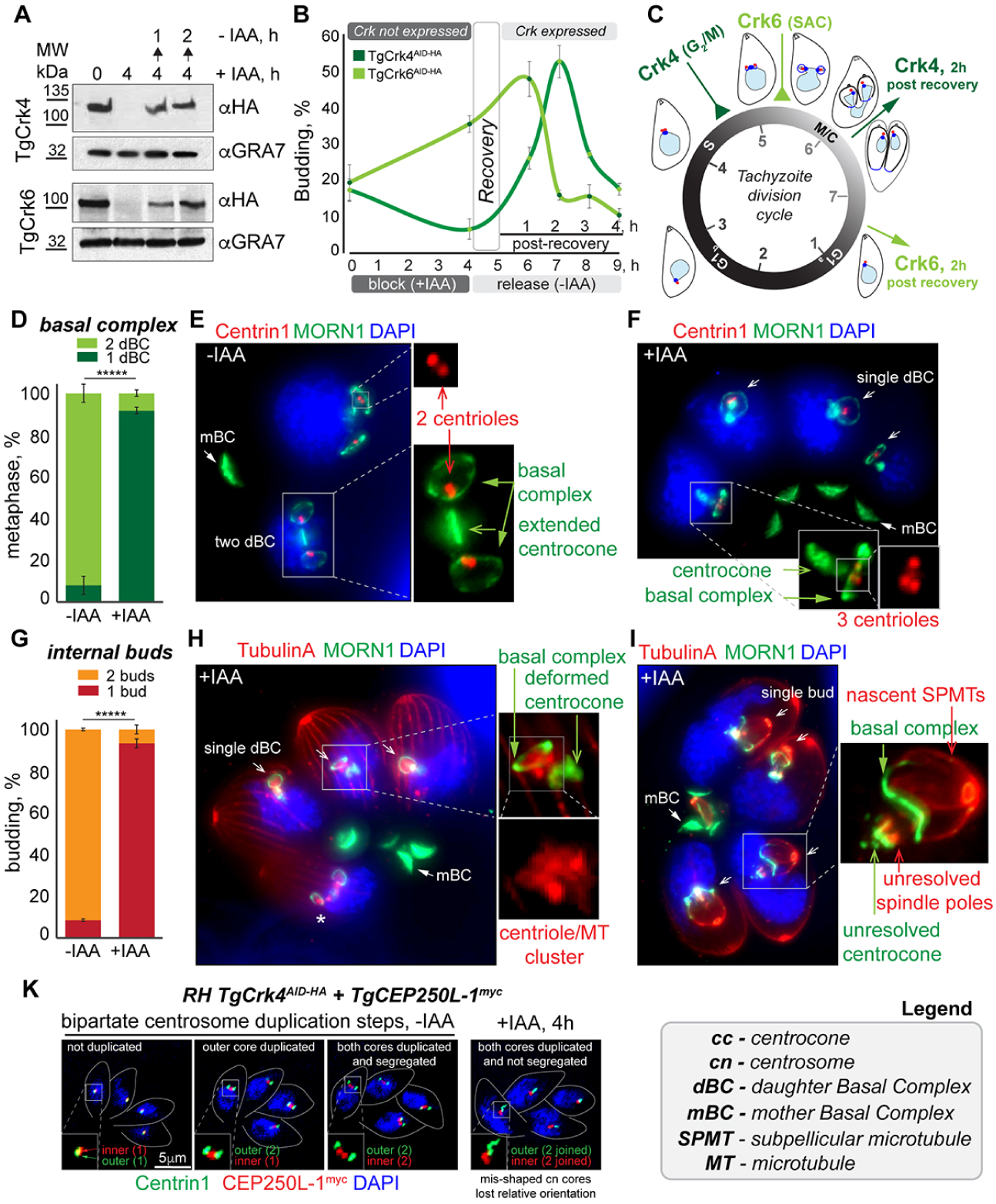
TgCrk4 regulates centrosome duplication and budding events upstream of the spindle assembly checkpoint. (A) Western Blot analysis of the total lysates of the RH*ΔKu80TIR1* tachyzoites expressing TgCrk4^AID-HA^ (upper panels) or TgCrk6^AID-HA^ (bottom panels). Lysates of non-treated parasites, parasites treated with 500μM IAA for 4 hours and chased for 1 and 2 hours were analyzed. Western blots were probed with α-HA (α-rat IgG-HRP) and with α-GRA7 (α-mouse IgG-HRP) to confirm equal loading of the total lysates. (B) Quantification of the budding populations of TgCrk4^AID-HA^ (dark green) or TgCrk6^AID-HA^ (light green) expressing tachyzoites during block (4 hours with 500μM IAA) and release (5 hours followed IAA removal). Recovery period marks first hour after IAA was removed from the growth medium. The mean -/+ SD values of three independent experiments are plotted on the graph. The t-test values are listed in TableS4. (C) The cell cycle diagram indicates relative positions of TgCrk6-and TgCrk4-regulated blocks at SAC and G_2_/M checkpoints and the timing of a 2-hour recovery from each block calculated from semi-synchronization experiments shown in panel B. (D) Quantification of the primary defect caused by RH TgCrk4^AID-HA^ deficiency. The centrocones connected to 2 (2dBC: light green) or 1 daughter basal complex (1dBC: dark green) were quantified in non-treated and treated with 500μM IAA for 4 hours parasites using TgMORN1 staining. A hundred random vacuoles of the parasites were examined in three independent experiments. The mean values are plotted on the graph. The t-test values are listed in TableS4. (E and F) The ultra-expansion microscopy analysis of RH*ΔKu80TIR1* TgCrk4^AID-HA^ expressing (E) and deficient (F) tachyzoites. Panel E: Co-staining of Centrin1 (α-Centrin1/α-mouse IgG Fluor 568) and TgMORN1 (α-MORN1/α-rabbit IgG Flour 488) shows duplicated centrosomes containing 2 centrioles, extended centrocone and two daughter basal complexes (dBC) organization in the metaphase. TgMORN1 localization in the mother basal complex (mBC) is indicated. Nucleus stained with DAPI (blue). Panel F: co-staining of the TgCrk4 deficient parasites (4 h, 500μM IAA) revealed a single dBC with multiple centrioles connected to deformed centrocone. (G) Quantification of the budding defect caused by RH TgCrk4^AID-HA^ deficiency. The number of the buds per cell were quantified in non-treated and treated with 500μM IAA for 8 hours using TgIMC1 staining. The longer IAA treatment allowed development of the bigger buds to aid quantifications. A hundred random vacuoles of the parasites were examined in three independent experiments. The mean values are plotted on the graph. The t-test values are listed in TableS4. (H and I) The ultra-expansion microscopy analysis of RH*ΔKu80TIR1* TgCrk4^AID-HA^ deficient tachyzoites. Staining with Tubulin A (α-TubulinA/α-mouse IgG Fluor 568) depicts subpellicular, centriolar microtubules (MTs) and the lack of spindle MTs. The TgMORN1 (α-MORN1/α-rabbit IgG Flour 488) staining shows changed morphology of the centrocone and number of the daughter basal complexes (dBC). Nucleus stained with DAPI (blue). (K) Immunofluorescent microscopy analysis of RH*ΔKu80TIR1* tachyzoites co-expressing TgCrk4^AID-HA^ and TgCEP250-L1^myc^. Parasites were co-stained with α-Centrin1 (α-mouse IgG Fluor 488) and α-myc (α-rabbit IgG Flour 568) antibodies to visualize the outer (Centrin1) and inner (TgCEP250 L1) cores of the bipartite centrosome. Three images of non-treated tachyzoites (-IAA) show the normal duplication of the centrosomal cores and morphological changes caused by TgCrk4 deficiency (+IAA, 4 hours) are shown on the right panel.

We treated TgCrk4 and TgCrk6 AID parasites with auxin for 4 hours (block) and monitored the release from block over 5 hours. Our results showed that retention of asynchronously dividing tachyzoites at a checkpoint for half of a division cycle led to significant enrichment of a specific cell cycle population (Fig. 2B). To determine the timing of TgCrk4-dependent arrest, we quantified parasites progressing through the budding stage. The asynchronous population of tachyzoites contained around 20% of parasites undergoing budding, which was detected with the budding marker TgIMC1. Contrary to TgCrk6-dependent arrest that led to a twofold increase, TgCrk4-dependent arrest reduced the number of budding parasites. Quantification showed that less than 10% TgCrk4-deficient parasites had internal buds, suggesting that TgCrk4 block occurs earlier than the TgCrk6 arrest. After TgCrk4 and TgCrk6 activities were restored (within one hour), the arrested parasites progressed to the next stage of the cell cycle in near synchrony (Fig. 2B). According to the budding dynamics, our semi-synchronization approach resulted in a 10-fold enrichment of cell cycle populations. Quantifying budding parasites upon release from block confirmed the difference in the timing of TgCrk4-and TgCrk6-regulated processes. Tachyzoites with restored TgCrk4 activity reached the budding peak an hour later than parasites with restored TgCrk6 expression (Fig. 2B and C). Based on our findings, we placed the TgCrk4-dependent process approximately 1 hour (1/7 of cell division duration) upstream of the SAC that controls metaphase-to-anaphase transition in mitosis. The relative positions of these regulatory steps suggests that TgCrk4 is an ideal candidate to regulate entry into mitosis at the G_2_ period that was presumed missing in apicomplexan cell cycles.

### Knockdown of TgCrk4 prevents segregation of duplicated centrosomes and spindle assembly

To determine what processes are regulated by the TgCrk4/TgCyc4 complex, we examined the phenotypes of tachyzoites lacking subunits of this complex. Since downregulation of TgCyc4 in the tet-OFF model was too lengthy to isolate the primary phenotype, we focused on TgCrk4 deficiency that can be achieved within 10 min of auxin treatment of TgCrk4 AID parasites. Given that the G_2_ period was considered missing in apicomplexans, there are no stage-specific markers. Therefore, we used a combination of Centrin1, TgMORN1, Tubulin A, and TgIMC1 markers to resolve the neighboring S-phase and mitosis concurrent with budding [11, 16, 18, 19]. Examination of TgMORN1-positive structures by ultra-expansion microscopy (U-ExM) showed that over 90% of TgCrk4-deficient parasites had altered the ratio of centrocone-to-daughter basal complexes (dBC). Normal progression throughout tachyzoite S-phase and mitosis involves the expansion of the intranuclear compartment centrocone, followed by development of two dBCs either attached or near the centrocone [18]. As the parasite transitions to anaphase in mitosis, the spindle, which is assembled in the centrocone, breaks, leading to centrocone resolution into two compartments, each of which is associated with a single dBC [11, 18]. We found that, instead of the typical two dBCs associated with the single centrocone, TgCrk4-depleted parasites had a single dBC linked to a deformed centrocone (Fig. 2D, E and F). Although it reminisced the ratio seen in anaphase, TgCrk4-deficient tachyzoites had a single unresolved centrocone linked to a single dBC (Fig. 2H, asterisk indicates normal anaphase). TgCrk4 deficiency also affected apicoplast segregation and fission that occurs concurrently with mitosis (Fig. S2B and C). Co-staining of the centrocone and dBC (MORN1) with microtubules (Tubulin A) or with the inner membrane complex (TgIMC1) confirmed the reduction of daughter cytoskeletal structures (Fig. 2H and I; Fig. S3). Nearly all budding parasites began assembling a single internal bud templated on a single dBC (Fig. 2G, H and I, Fig. S2).

The appearance of a single centrocone linked to a single dBC can result from a failed centrosome duplication or duplicated centrosomes that failed to segregate. Our observations strongly support the latter. Staining with α-Tubulin A and α-Centrin1 antibodies revealed centrioles and possibly short microtubule (MT) fibers accumulated near the unresolved centrocone (Fig. 2F and G). The presence of more than 2 centrioles in the cluster confirmed that TgCrk4-deficient parasites can successfully duplicate centrosomes. The clustered centrioles near a single centrocone confirmed that duplicated centrosomes did not segregate even in budding parasites. The *T. gondii* centrosome is composed of inner and outer cores with differential protein composition [15, 16, 20]. TgCentrin1 is localized at the outer, centriole-containing core while TgCEP250-L1 is preferentially expressed in the inner centrosomal core [19]. To determine if TgCrk4 downregulation affects the inner core, we placed a 3xmyc epitope tag on endogenous TgCEP250-L1 protein in the RH TgCrk4^AID-HA^ transgenic line (Fig. 2K). IFA analysis of TgCrk4-expressing tachyzoites confirmed the expected stepwise duplication of both outer and inner cores of the centrosome. Examination of tachyzoites treated with 500μM IAA for 4 hours revelated that centrosomal cores were affected by TgCrk4 deficiency. Judging by their size and morphology, both cores appeared to be duplicated but failed to segregate. In addition, the centrosomal cores lost their proper orientation relative to one another, often placed orthogonally rather than parallel to each other.

The inner core of the centrosome is tightly aligned with the centrocone [16]. In the normal *T. gondii* mitosis, centrocone extension coincides with spindle pole growth (Fig. 2E and Fig. S3A) [11, 16, 18]. Examination of TgCrk4-deficient parasites revealed that deformed centrocones did not incorporate an extended pole-like structure, implying that TgCrk4-deficient parasites fail to assemble a bipolar spindle (Fig. 2H and I, Fig. S3B). Thus, we concluded that TgCrk4-deficient parasites could neither segregate duplicated centrosomes nor form a bipolar spindle. In the conventional cell cycle, segregation of duplicated centrosomes is a prerequisite for bipolar spindle assembly, which signifies the entry into mitosis, also known as the G_2_/M transition [21]. The similar regulation of these processes and the spatiotemporal localization of the TgCrk4-TgCyc4 complex suggest that, contrary to previous assumptions, the G2 period does operate in apicomplexan cell cycles. In *T. gondii* endodyogeny, the G_2_/M transition is regulated by the novel TgCrk4-TgCyc4 complex.

### TgCrk4 represses multinuclear division by blocking centrosome and chromosome reduplication

To further verify the role of TgCrk4 in centrosome segregation and spindle assembly, we examined the recovery of TgCrk4-deficient tachyzoites during the first three hours of restored TgCrk4 expression (Fig. 3). In good agreement with the predicted role of TgCrk4 in centrosome segregation, tachyzoites released from TgCrk4-dependent block began to separate their centrosomes. The number of separated Centrin1-positive centrosomes increased from 10% to 50% (Fig. 3A, B, C and D). Surprisingly, we noticed the emergence of parasites with three or more centrosomes concurrently to an increase of parasites with multiple internal buds (Fig. 3D and E).

**Figure 3.**
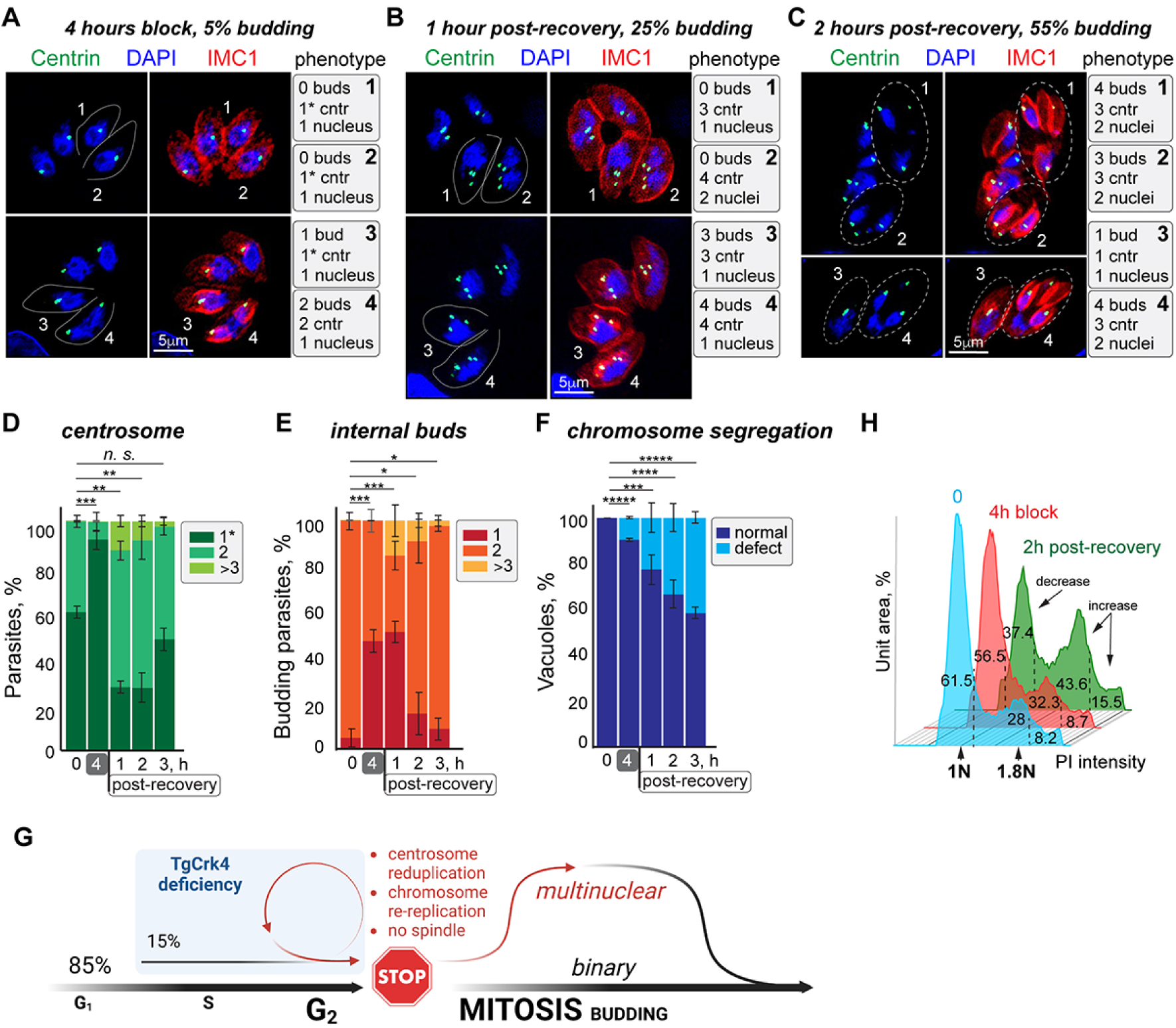
TgCrk4 deficiency leads to centrosome reduplication and DNA rereplication. (A, B and C) Immunofluorescent microscopy analysis of RH*ΔKu80TIR1* TgCrk4^AID-HA^ tachyzoites incubated with 500μM IAA for 4 hours (A) [block] and then without IAA for 2 (B) [1-hour post-recovery] or 3 (C) [2 hours post-recovery] hours. Parasites were co-stained with α-Centrin1 (α-mouse IgG Fluor 488), α-TgIMC1 (α-rabbit IgG Fluor 488) antibodies and DAPI to estimate the ratio of centrosome, nuclei and internal buds per parasite. Analyzed cells are numbered and the summary is shown on the right of each panel. (D, E and F) Number of centrosomes (D) and internal buds (E) per parasite, and DNA mis-segregation defects (F) were quantified in asynchronously growing (0 hours) RH*ΔKu80TIR1* TgCrk4^AID-HA^ tachyzoites as well as in the arrested (4 hours with 500μM IAA, grey block), and recovered (1, 2 and 3 hours after kinase expression was restored) populations. A hundred random vacuoles of the parasites were examined in three independent experiments. The mean values are plotted on the graphs. The t-test values are listed in TableS4. (H) FACScan analysis of DNA content of TgCrk4^AID-HA^ expressing (0, blue plot), TgCrk4^AID-HA^ deficient (4 hours with 500μM IAA, red plot) and recovered from TgCrk4^AID-HA^ deficiency (3 hours after IAA removal, green plot) parasites. Dashed lines show the gates for non-replicated (<1-1.2N), replicated (1.2-2N) and over-replicated (>2N) DNA. (G) Cell cycle effect of the TgCrk4 deficiency. The schematics shows the cell cycle progression of TgCdt1 deficient tachyzoites. Most of the parasites quickly recover from TgCrk4-dependent G_2_ arrest (red STOP sign). About 15% of the tachyzoites that spent a longer time in the block re-enter S-phase (blue block: centrosome and chromosome reduplication), which led to one-cycle multinuclear division. The surviving DNA-mis-segregation defect population quickly reverted to binary division upon restoration of TgCrk4 activity.

FACScan analysis of their DNA content confirmed that release from block led to decrease of G_1_ and concurrent increase of populations that duplicated and over-duplicated DNA (Fig. 3H). During 2h of post-block recovery, the number of parasites whose DNA content exceeded 2N doubled, suggesting that a fraction of TgCrk4-deficient parasites reduplicated their chromosomes and centrosomes. On the contrary, DNA mis-segregation had amplified during recovery, confirming that it was a secondary effect of TgCrk4 deficiency, in particular, the outcome of centrosome reduplication (Fig. 3F). The phenotypic analysis of parasites emerging from TgCrk4-induced block showed that TgCrk4-deficient parasites broke the “once per cell cycle” rule of DNA and centrosome duplication [22, 23].

In the conventional cell cycle, centrosome duplication is tightly linked to chromosome replication [24]. Centrosome duplication happens at the G_1_/S transition and is blocked for the rest of the division cycle [22]. Likewise, licensing of DNA replication takes place in G_1_ and is forbidden in S and G_2_ phases [25]. The major goal of G_2_ period is to repress an unlawful DNA rereplication and centrosome reduplication [26]. The fact that TgCrk4-deficient parasites had multiple centrosomes and over-replicated chromosomes indicated that S-phase events were reinitiated, which happens only in multinuclear divisions such as those employed by apicomplexan schizogony. The fraction of parasites with multiple centrosomes or buds likely represented a population that spent the longest time with arrested cell cycles (Fig. 3G). It is possible that non-segregated centrosomes serve as a signal of incomplete DNA replication, allowing relicensing and refiring of DNA replication origins, and reinitiating centrosome duplication. The quick reduction of the number of multinuclear dividing parasites released from the TgCrk4-dependent block further confirmed the primary role of TgCrk4 in the repression of multinuclear and promotion of binary division. The accompanied growth of mis-segregation defects suggests that the switch from binary to multinuclear division involves multiple regulators, some of which may not be expressed in tachyzoites programmed for binary division.

### Global profiling of TgCrk4-induced cell cycle block

To determine the cellular effect of TgCrk4 deprivation, we examined changes in global protein expression and phosphorylation in tachyzoites that were held in a TgCrk4-induced block for 4 hours (Fig. 4A and B). Our results showed specific changes in the protein landscape that corroborated our immunofluorescent microscopy analyses of TgCrk4-deficient tachyzoites (Fig. 4A, Fig.2 and Fig.3). We detected a nearly complete tachyzoite proteome (4219 proteins), out of which the TgCrk4-deprived tachyzoites had 58 upregulated and 126 downregulated proteins (Table S3). In good agreement with the predicted role of TgCrk4 in repressing chromosome rereplication, GO term analysis detected upregulation of DNA replication licensing factors TgORC4, TgORC5, and DNA polymerase. Since DNA repair is a process that takes place in G_2_ phase, the increase in expression of DNA repair proteins in TgCrk4-deficient tachyzoites further confirms the role of TgCrk4 as a G_2_ phase regulator. Corroborating TgCrk4-dependent suppression of budding, we detected decreased expression of inner membrane complex proteins that are enriched on daughter buds, such as IMC16, IMC30, IMC33 and IMC34 [27–29]. Regulated proteolysis plays a vital role in the progression through S, G_2_ phase, and mitosis [30]. The TgCrk4-induced block led to the downregulation of deubiquitinase TgOTUD3A and caspase TgMCA1. Interestingly, previous studies by Dr. Sinai’s group showed that TgOTUD3A knockdown induced a switch from binary to multinuclear division, which resembles the TgCrk4-deficiency phenotype and suggests a functional link between these cell cycle factors [31].

**Figure 4.**
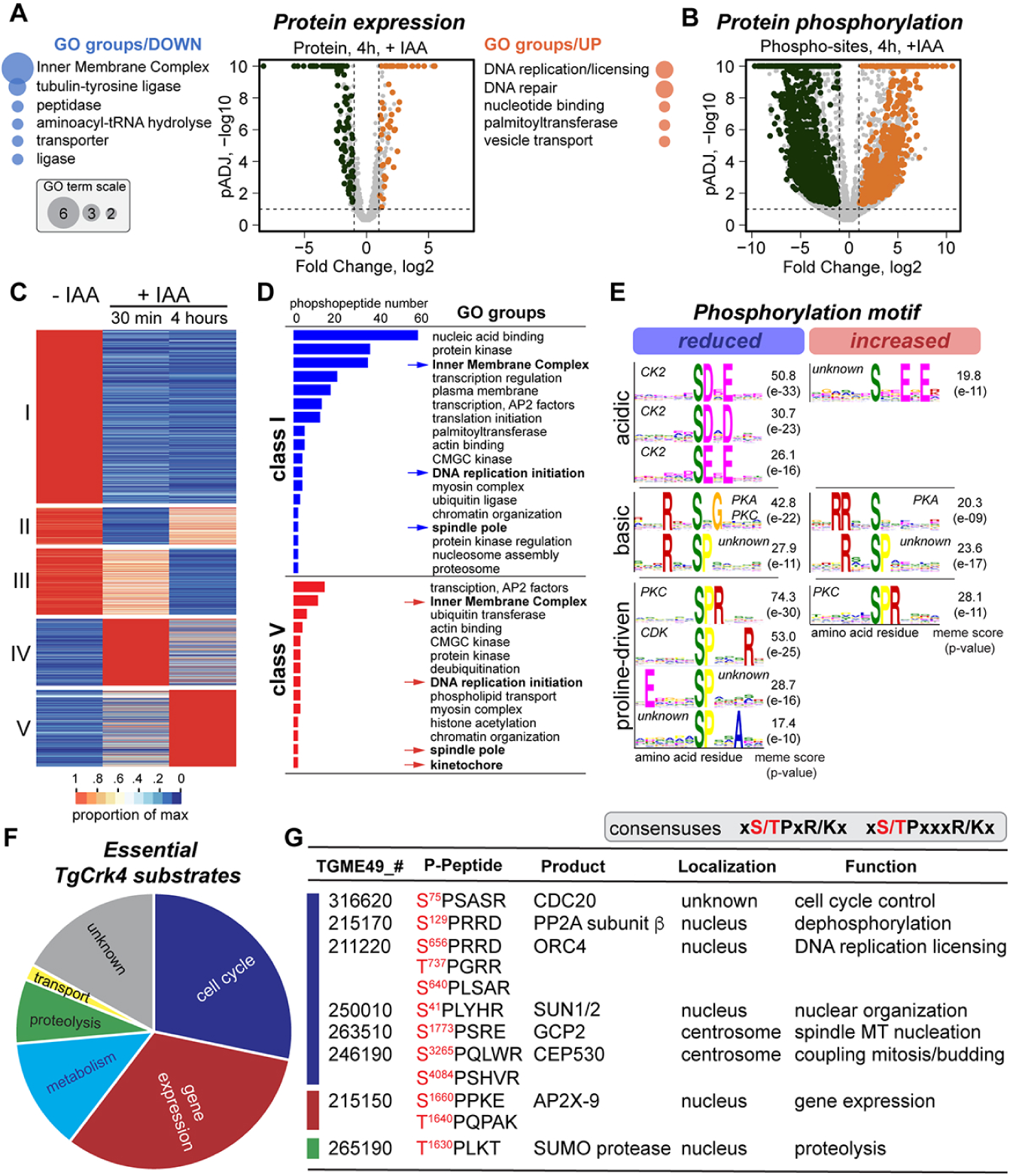
Global proteomic and phosphoproteomic analysis of *T. gondii* G_2_/M checkpoint. (A and B) Volcano plots show changes of the protein expression level (A) and of the phospho-sites intensity (B) at the checkpoint block induced by TgCrk4^AID-HA^ downregulation 4 hours. The GO term enrichment analysis performed on upregulated (orange) and downregulated (blue) groups identified in global proteome. Results are shown on the sides of each plot. The bubbles reflect the size of individual pools. (C) The heat map displays global changes of the protein phosphorylation caused by auxin induced TgCrk4^AID-HA^ degradation for 30 min and 4 hours. The proteins are organized according to the similarity in phosphorylation by K-means and combined into five classes based on the temporal dynamic of phosphorylation. (D) The graphic representation of the GO term analysis of class I (reduced phosphorylation) and V (increased phosphorylation) phospho-peptides. The groups of particular interest are indicated with arrows and shown in bold. (E) The phosphorylation motifs affected by 30 min TgCrk4^AID-HA^ degradation were deduced using momo software. Three-letter motifs with p-values >e-10 are shown. Responsible kinase family and scores are indicated on the corresponding plots. (F) Pie chart of the putative TgCrk4 substrate proteins. The essential proteins that reduced or lost intensity of TgCrk4-dependent phosphorylation were categorized. (G) A selected set of putative TgCrk4 substrates that were identified based on the reduction of phosphorylation intensity within a Proline-driven motif (consensus) caused by the lack of TgCrk4 for 30 min and 4 hours (Class I, panel C). Affected phospho-Serine or phospho-Threonine residues are shown in red.

The comparative global phosphoproteome of asynchronous and TgCrk4-arrested tachyzoites contained 23,480 phospho-peptides (Table S3). TgCrk4-dependent cell cycle arrest resulted in increased phosphorylation of 1112 and decreased phosphorylation of 883 peptides (Fig. 4B). Examining short-(30 min) and long-term (4 hours) TgCrk4 deprivation revealed five patterns (classes) of temporal phosphorylation (Fig. 4C). We performed a GO term enrichment analysis of class I and V proteins that showed the loss or steady increase of phosphorylation, respectively. TgCrk4 knockdown significantly affected the protein expression machinery that controls transcription, translation, and regulated proteolysis. A dominating group of nucleic acid binding proteins included 16 AP2 DNA binding factors, among which are three transcriptional repressors AP2XII-1, AP2XI-2, and AP2IX-9, which function is linked to parasite development [32, 33]. A few histone modifiers and nucleosome assembly factors also appeared to be regulated by alternative phosphorylation at the G_2_ block. Protein kinases constituted the second large group affected by TgCrk4 downregulation and included essential for tachyzoite replication kinases, MAPK1 and TKL2 [16, 34, 35]. We had previously shown that activating a temperature-sensitive allele of TgMAPK1 led to centrosome amplification, which placed TgMAPK1 into the broader G_2_ network [16]. We detected 34 IMC proteins that lost and 11 that gained phospho-modifications, which likely reflects the repressed budding in TgCrk4-deficient tachyzoites. In good agreement with TgCrk4 role in regulating G_2_, both class I and class V proteins included DNA replication licensing factors MCM2, 4, 6, and 7, phosphorylation of which may be a part of the mechanism that represses relicensing of DNA replication. Furthermore, TgCrk4 depletion resulted in the altered phosphorylation of γ-tubulin ring complex proteins GCP2 and GCP4 that nucleate spindle MTs, and increased phosphorylation of the apicomplexan-specific kinetochore proteins AKIT1 and AKIT6, which suggests they may regulate repression of the spindle and kinetochore assembly [36].

To find out what kinases operate in the *T. gondii* G_2_ phase, we determined phosphorylation motifs that were affected in the brief absence of TgCrk4 (30 min) (Fig. 4E). The MEME analysis revealed a specific reduction of phosphorylation within 33 motifs and increased phosphorylation within 14 motifs representing three major groups: acidic, basic, and proline-driven motifs. The loss of TgCrk4 affected casein kinase 2 (CK2), Cdk-, and PKC/PKA-dependent phosphorylation. While the activity of PKC and PKA seem to be differentially modulated, CK2 and Cdk activities were specifically reduced in response to TgCrk4 removal. Interestingly, TgCrk4 degradation largely affected proteins containing an uncommonly extended Cdk motif [(S/T)PxxxK/R], adding another non-conventional feature to TgCrk4. In addition, we detected several unknown or highly deviated phosphorylation motifs that implicate the activity of novel kinases in the control of G_2_ phase.

### Search for TgCrk4 substrates

To identify immediate TgCrk4 effectors, we interrogated phospho-peptides that contain the most common (S/T)*PxR/K and extended (S/T)*PxxxR/K Cdk motifs which had lost or reduced phosphorylation both immediately (30 min, +IAA) and after prolonged TgCrk4 downregulation (4 hours, +IAA) (Fig. 4E) [37]. Out of 109 peptides that matched the criteria, 54 were predicted to be essential proteins (Table S3). Based on ToxoDB database annotations, the most abundant categories of putative TgCrk4 substrates were proteins involved in cell cycle regulation (15 peptides from 12 proteins) and gene expression control (17 peptides from 15 proteins) (Fig. 4F). Validating the role of TgCrk4 in chromosome rereplication, three residues of the DNA replication licensing factor TgORC4 (TGME49_211220), S^656^, S^737^ and T^737^ were predicted to be dependent on TgCrk4 phosphorylation (Fig. 4G). Coincidently, global protein expression analysis of TgCrk4-deficient tachyzoites showed increased TgORC4 expression (Fig. 4A). This suggests that one of the mechanisms controlling chromosome rereplication in G_2_ phase could be affected by reduced stability of phosphorylated TgORC4. The list of putative substrates included the repressor of the anaphase-promoting complex/cyclosome (APC/C) Cdc20 (TGME49_316620) (Fig. 4G). In conventional cell cycle, the Cdc20 phosphorylation occurs at the G_2_/M transition, which further confirms that TgCrk4 regulates G_2_ phase in tachyzoites [38]. Furthermore, our results suggest that TgCrk4 likely controls initiation of spindle pole assembly by directly phosphorylating TgGCP2 (TGME49_263510), a component of the γ-tubulin ring complex responsible for nucleating spindle microtubules [39]. Interestingly, we also detected factors that were not previously implicated in G_2_ phase control, such as the nuclear membrane protein SUN1/2 (TGME49_250010), bipartite centrosome protein TgCEP530 (TGME49_246190) and DNA binding factor TgAP2X-9 (TGME49_215150), suggesting novel routes of G_2_ regulation [20] (Fig. 4G). It was recently shown that TgAP2X-9 transcription is directly regulated by another AP2 factor, TgAP2IX-5 implicated in the control of binary division in *T. gondii* tachyzoites [40]. Lastly, the progression through the G_2_/M transition involved specific changes in protein expression and phosphorylation that could be dependent on TgCrk4 phosphorylating the TgSUMO protease (TGME49_265190) and a regulatory β-subunit of TgPP2A phosphatase (TGME49_215170).

### Functional orthologs of TgCrk4-TgCyc4 complex are only present in binary dividing apicomplexans

Previous phylogenetic analyses of apicomplexan Cdk-related kinases showed that Crk4 orthologs exist in apicomplexans and ancestral alveolates [6]. We performed a follow-up examination of the latest annotated genomes and confirmed the relatively broad inheritance of Crk4. TgCrk4 orthologs were found in three core Apicomplexa groups, coccidians, piroplasmids, and cryptosporidians, as well as in chrompodelids closest to the Apicomplexa lineage, which suggests that Crk4 kinase was inherited from apicomplexan ancestors (Fig. 5A). Contrary to its partner kinase, TgCyc4 had a limited presence in the analyzed genomes. Given the lack of detectable TgCyc4 orthologs in chrompodelids, it is likely that apicomplexans evolved rather than inherited this cyclin. The fact that TgCyc4-related factors were found only in several coccidian species and in piroplasmids further corroborated the possibility of an independent rise of Cyc4 factor in apicomplexans.

**Figure 5.**
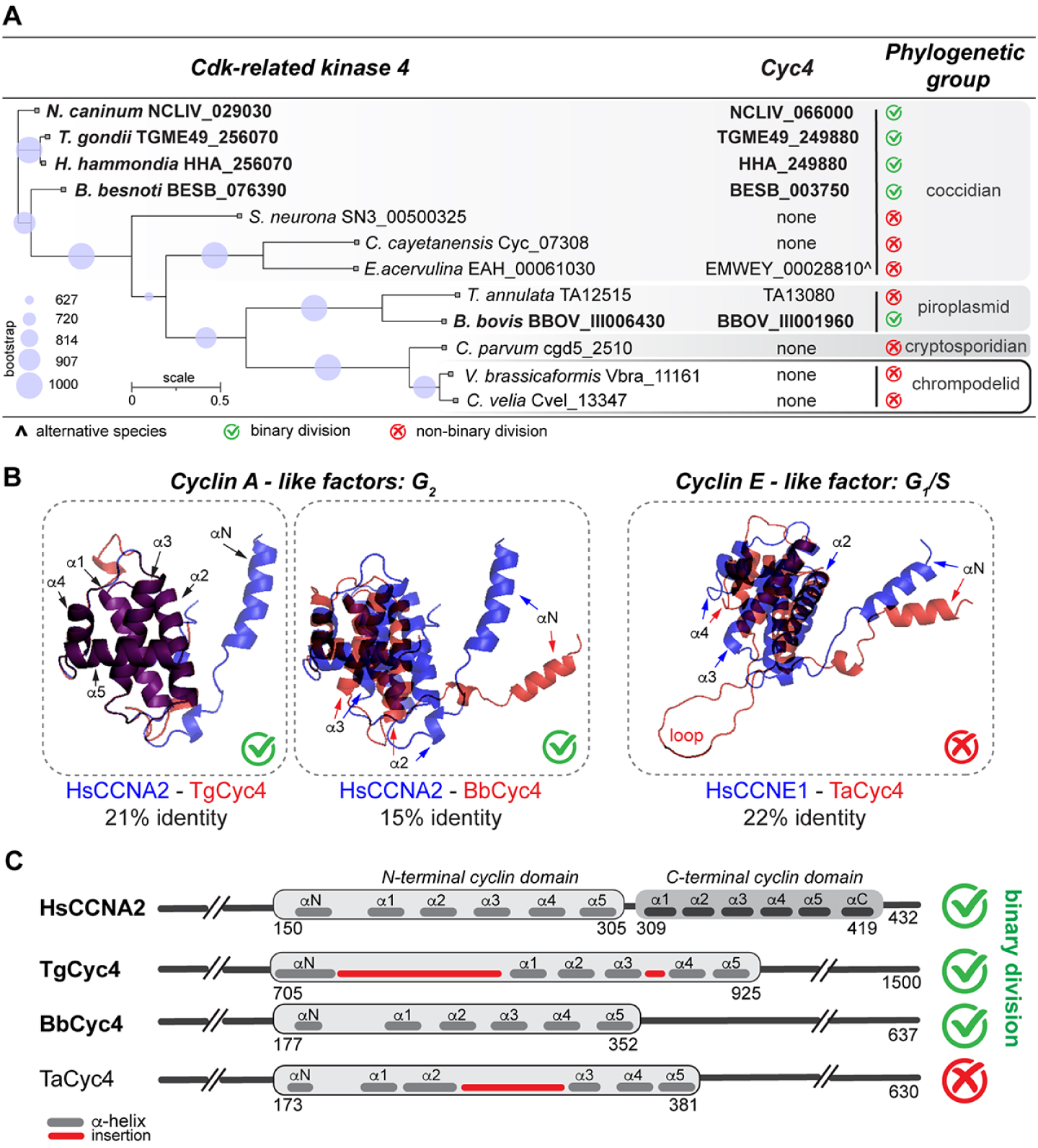
Structural and phylogenetic analysis of TgCrk4-TgCyc4 complex in apicomplexans. (A) Summary of the phylogenetic analyses of the TgCrk4 related kinases and the orthology search of TgCyc4 related factors among Apicomplexa groups and chromopodelids. (B) Folding prediction of the cyclin domains of *T. gondii* Cyc4, *Babesia bovis* Cyc4 and *Theileria annulate* Cyc4 overlayed with H. sapiens Cyclin A2 and Cyclin E1 based on best predicted fit (SWISS-MODEL). Changes in the structures are indicated with arrows. The percentage of identical amino acid residues in the modeled regions is shown below. (C) Comparative schematics of *H. sapiens* Cyclin A2 and TgCyc4, BbCyc4 and TaCyc4 protein organization.

Structural analyses of apicomplexan TgCyc4-related factors revealed a limited resemblance to mammalian Cyclin A and Cyclin E. Although highly similar at the level of amino acid residues, Cyclin A and Cyclin E regulate different cell cycle phases [41]. In mammalian cells, Cdk2-Cyclin E complex controls the entry into S-phase, while Cdk1/2-CyclinA complex is involved in regulating the G_2_/M transition. Interestingly, our modeling of apicomplexan Cyc4 revealed the resemblance of the coccidian and *Babesia* Cyc4 with G_2_ Cyclin A, while the *Theileria* Cyc4 was preferentially modelled to G_1_/S Cyclin E (Fig. 5B). It suggests that, while related, *Theileria* Cyc4 may regulate its cell cycle phase differently than how the phase is regulated by Cyc4 in other apicomplexans. Further examination of cyclin folds revealed the *Eimeria* ortholog of TgCyc4 had a significant alteration in its cyclin domain (Fig. S4). Cyclins are a highly divergent group of proteins with the unifying feature of two conservative cyclin domains located in the C-terminus. The cyclin domain typically responsible for binding and activation of Cdk type kinase is composed of six short α-helixes (Fig. 5C) [41]. Our AlphaFold2 analysis predicted drastic structural changes caused by sizable poly-Q insertions that fused α3, α4, and α5 helices, which would severely affect the *Eimeria* Cyc4 ability to bind or activate a Cdk kinase (Fig. S4B).

The coccidian Cyc4 factors found in *T. gondii*, *Hammondia hammondia*, *Neospora caninum,* and *Besnoitia besnoiti* also contained an insertion between N-terminal activating and α1 helix (Fig. 5B and C). However, our study of TgCrk4-TgCyc4 complex verified that the loop in TgCyc4 had no effect on TgCyc4 ability to selectively bind TgCrk4 (Fig. 1). Altogether, we identified six Apicomplexa genera that express functional TgCrk4-TgCyc4 related complex: *Toxoplasma*, *Hammondia*, *Neospora*, *Besnoitia*, *Babesia, and Theileria*. Structural similarity with G_2_ Cyclin A suggested that the Crk4-Cyc4 complex of *Toxoplasma*, *Hammondia*, *Neospora*, *Besnoitia*, and *Babesia* may control G_2_ events. On the contrary, similarity of *Theileria* Cyc4 and Cyclin E indicated the involvement of *Theileria* Crk4-Cyc4 complex in regulation of the different cell cycle stages or different cellular processes. Coincidently, unlike most apicomplexans, *Toxoplasma*, *Hammondia*, *Neospora*, *Besnoitia*, and *Babesia* divide in binary fashion such as endodyogeny (*T. gondii*, *H. hammondia*, *N. caninum*) or binary fission (*Babesia* spp.) [3]. Based on our examination of TgCrk4-TgCyc4 function in *T. gondii* and our phylogenetic and structural analysis across Apicomplexa phylum, we propose that TgCrk4-TgCyc4 complex and its functional orthologs had evolved to repress multinuclear division in a subgroup of apicomplexan parasites.

### TgCrk4-TgCyc4 complex interacts with a Cdt1-like factor

The interactomes of TgCrk4 and TgCyc4 identified the TGME49_247040 protein with unknown function as a dominant interactor of the complex. Although TGME49_247040 protein does not have any recognizable domains, Swiss-Prot modeling detected a partial resemblance to the DNA replication licensing factor Cdt1 (Fig. 6A). Mammalian Cdt1 is composed of an N-terminal regulatory and C-terminal minimal origin licensing domain [42]. Besides its interaction with the inhibitor geminin and a mini-chromosome maintenance (MCM) replicating helicase complex, Cdt1 directly binds to DNA polymerase loading factor PCNA1 (PIP box) and G2 cyclin A (Cy motif) [42]. We found that TGME49_247040 had limited homology to the geminin-and MCM-binding regions of mammalian Cdt1. Further examination revealed that, like eukaryotic Cdt1, TGME49_247040 factor had several recognizable cyclin-binding motifs (Cy) and degrons (D-box). In addition, TGME49_247040 protein was predicted to have nuclear localization signals (NLS) (Fig. 6A). To find out if TGME49_247040 is orthologous to the Cdt1 proteins of other eukaryotes, we performed phylogenetic analyses of apicomplexan TGME49_247040-related proteins and Cdt1 orthologs from mammalians, land plants, fungi, and dinoflagellates (Fig. 6B). The results supported a common inheritance of apicomplexan and eukaryotic Cdt1 factors, and we named TGME49_247040 protein TgCdt1. Our results revealed that, contrary to predictions, apicomplexan parasites encode Cdt1 orthologs that appeared to escape previous detection due to low sequence similarity.

**Figure 6.**
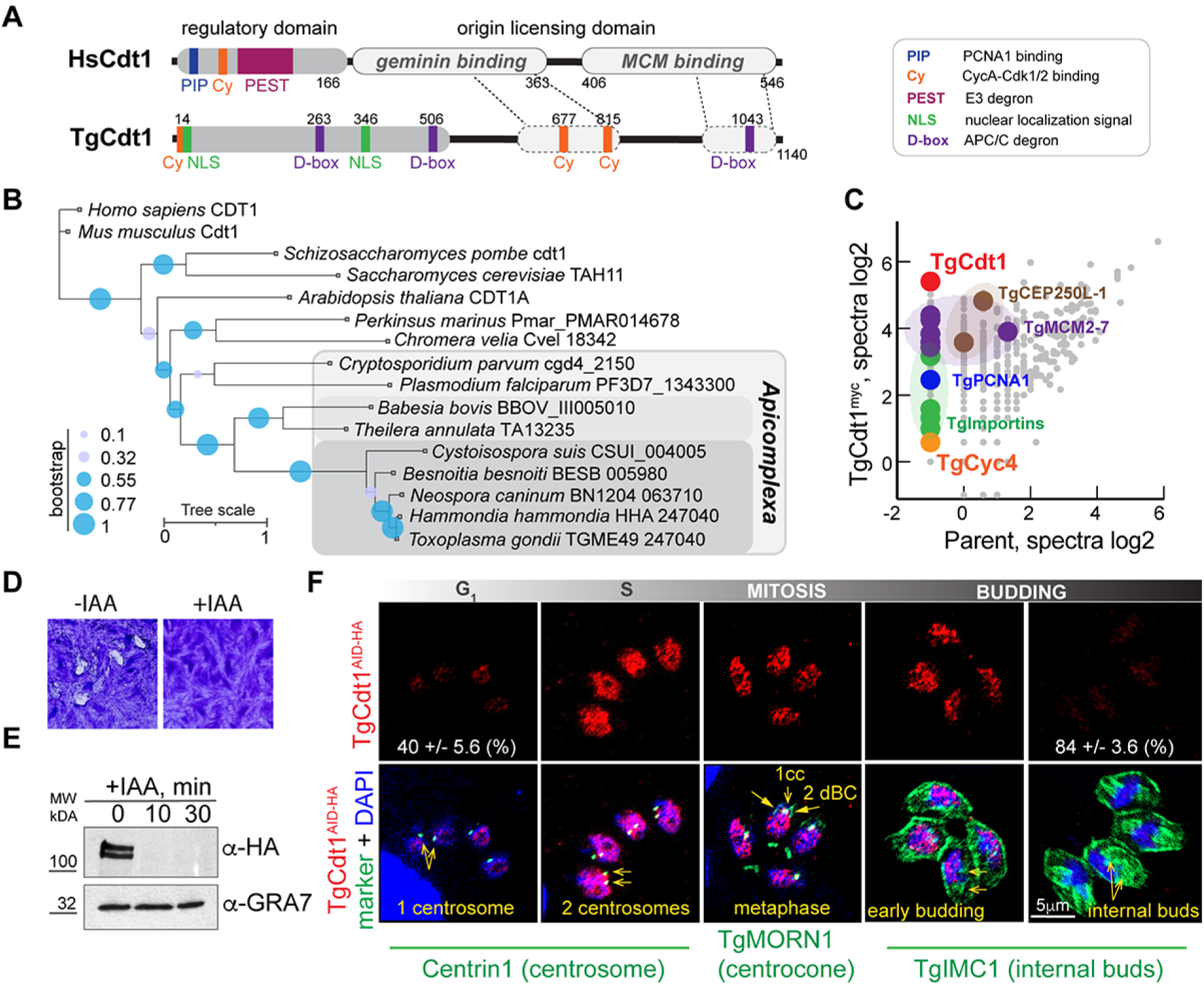
*T. gondii* expresses factor related to DNA replication licensing factor Cdt1. (A) Schematics of *H. sapiens* Cdt1 and *T. gondii* Cdt1 protein organization. The signature domains and regions of TgCdt1 are shown based on the ELM prediction. Dotted lines indicate two regions of similarity detected by SWISS-MODEL software. (B) Phylogenetic tree of apicomplexan Cdt1. (C) The log2 values of the protein spectra detected by mass-spectrometry analysis of the TgCdt1^myc^ complexes are plotted on the graph. The different color dots represent categories of the selected TgCdt1 interactors. (D) Images of the host cell monolayers infected with TgCdt1^AID-HA^ tachyzoites and grown with or without 500μM IAA for 7 days. (E) Western Blot analysis confirms downregulation of TgCdt1^AID-HA^ after 10 and 30-min treatment with 500μM IAA. Western blots were probed with α-HA and α-GRA7 to confirm equal loading of the total lysates. (F) Immunofluorescent microscopy analysis of TgCdt1^AID-HA^ cell cycle expression. The TgCdt1^AID-HA^ (α-HA/α-rat IgG Fluor 568) co-staining with Centrin1 (α-Centrin1/α-mouse IgG Fluor 488) was used to distinguish G_1_ (1 centrosome) and S-phase (2 centrosomes). The mitotic metaphase was detected by co-staining with TgMORN1(α-MORN1/α-rabbit IgG Flour 488). The budding parasites were identified by co-staining with alveolar protein TgIMC1(α-IMC1/α-rabbit IgG Fluor 488). The blue staining represents nucleus (DAPI). Cell cycle phases were determined based on the number of the reference structures and morphology of the nucleus as indicated.

To confirm the interaction of TgCdt1 with the TgCrk4-TgCyc4 complex, we fused endogenous TgCdt1 with a 3xmyc epitope and isolated TgCdt1^myc^ complexes by affinity purification. In good agreement with the presence of Cy motifs, we detected a weak but specific interaction between TgCdt1 and TgCyc4 (Fig. 6C, Table S3). No other cyclins or cell cycle Crks were detected in the TgCdt1 proteome, suggesting that TgCdt1 binds TgCrk4-TgCyc4 complex via the TgCyc4 subunit. Studies in metazoans demonstrated a dominant Cdt1-geminin interaction [43]. However, despite containing a conserved geminin binding region, we could not identify a possible *Toxoplasma* ortholog or analog of the Cdt1 inhibitor, geminin. The absence of a geminin-like factor in TgCdt1 proteome suggests that, like budding yeast, *T. gondii* may not encode geminin [44].

### TgCdt1 controls DNA and centrosome reduplication in tachyzoites

Studies in metazoans and fungi established the role of Cdt1 in licensing DNA replication (late mitosis and G_1_), controlling chromosome rereplication (S-phase and G_2_), and segregation (mitosis) [42, 45–47]. Our TgCdt1 interactome showed the enrichment of conventional Cdt1 interactors such as a replisome complex (MCM2-7) and DNA polymerase loading factor TgPCNA1, pointing toward a conservative role of TgCdt1 in regulating DNA replication in *T. gondii* (Fig. 6C, TableS3) [48]. To examine the TgCdt1 function in tachyzoites, we built and analyzed a TgCdt1 AID conditional expression model. Immunofluorescence microscopy analyses confirmed the nuclear localization of TgCdt1^AID-HA^, which was consistent with predicted NLS motifs and TgCdt1 interactions with α-and β-Importins (Fig. 6C and F). Western blot analysis demonstrated the fast auxin-induced degradation of TgCdt1^AID-HA^, and a plaque assay verified that TgCdt1 was required for tachyzoite growth (Fig. 6D and E). Detailed examination of TgCdt1 function in tachyzoites revealed significant differences from the conventional eukaryotic Cdt1 factor. As a critical DNA replication licensing factor, metazoan and fungal Cdt1 is active on chromosomes in late mitosis and G_1_ phase [42]. To avoid unlawful rereplication of the chromosomes, Cdt1 dissociates from replicating chromosomes in S-phase, which is achieved by proteolysis of Cdt1 in metazoans, or by shuttling Cdt1 from the nucleus in budding yeast [42]. We found that, contrary to conventional Cdt1, TgCdt1 was abundantly expressed in the nucleus of S-phase tachyzoites and was absent in late mitosis and early G_1_ phase (Fig. 6F).

The unpredicted spatiotemporal expression raised doubt over if TgCdt1 functions as a DNA replication licensing factor in *T. gondii*. To address this question, we examined the DNA content in tachyzoites either expressing or depleted of TgCdt1 for the duration of one division cycle (8h). Our FACScan analysis of TgCdt1 AID tachyzoites did not detect the anticipated repression of DNA replication in TgCdt1-deficient tachyzoites (Fig. 7A). On the contrary, eliminating TgCdt1 led to a decrease of the G_1_ population (1N DNA content) and was accompanied by an increase in parasites both undergoing DNA replication (1.8 DNA content) and with re-replicated DNA (>2N DNA content). Quantitative IFA of TgCdt1-deficient tachyzoites verified the substantial reduction of parasites in G_1_ phase and the expansion of replicating (S-phase) and mitotic populations (Fig. 7B). These findings strongly suggested TgCdt1 has minor to no involvement in licensing DNA replication in tachyzoites.

**Figure 7.**
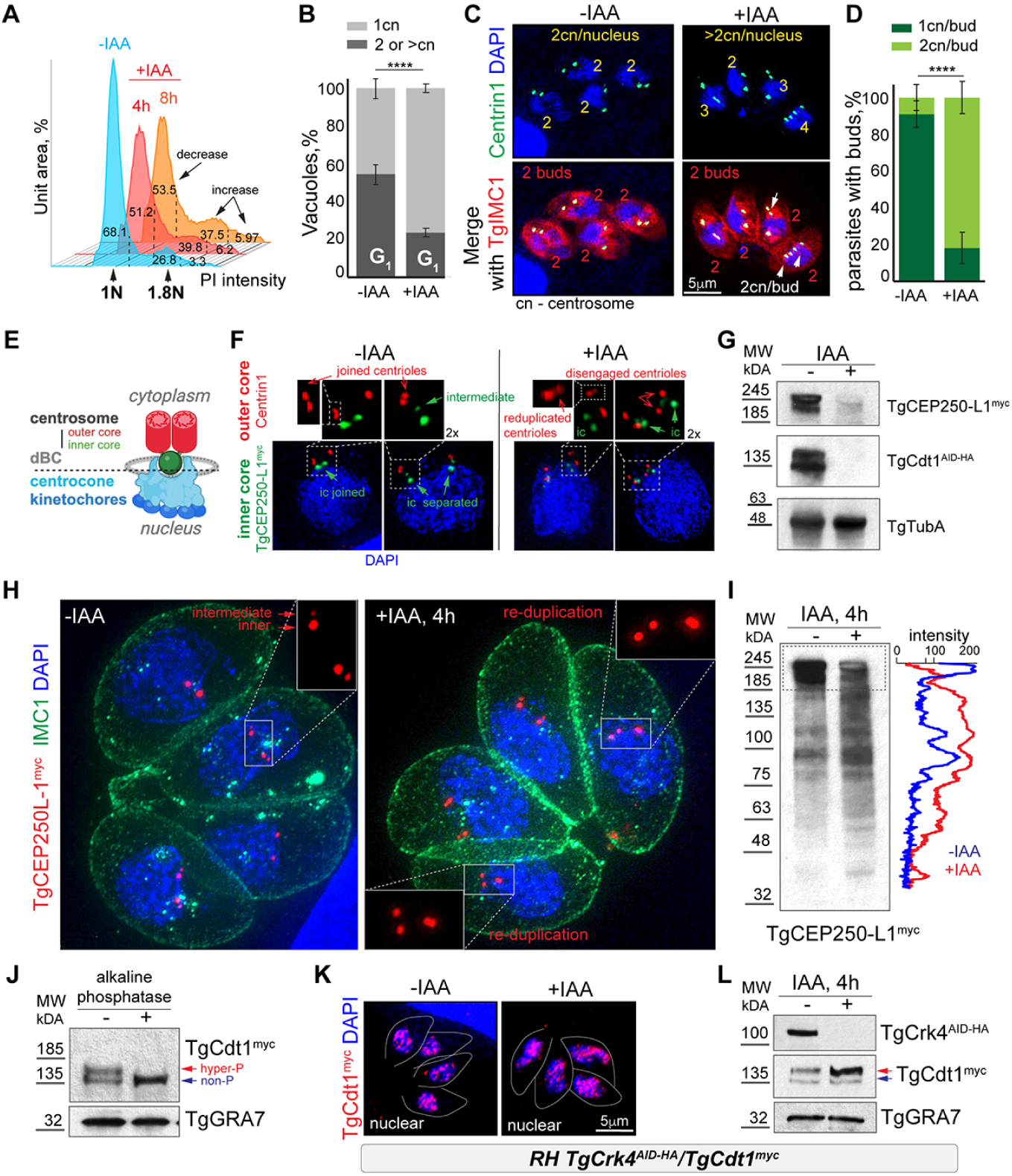
TgCdt1 is a phosphorylated protein that controls centrosome reduplication and chromosome rereplication. (A) FACScan analysis of DNA content of TgCdt1^AID-HA^ expressing (0, blue plot) and TgCdt1 deficient for 4 hours (+IAA, red plot) and 8 hours (+IAA, orange plot) parasites. Dashed lines show the gates used to segregate parasites containing non-replicated (<1-1.2N), replicated (1.2-2N) and over-replicated (>2N) DNA. (B) Quantification of the G_1_ (single centrosome) and S/G_2_/M/budding (2 centrosomes) populations. The TgCdt1^AID-HA^ expressing and TgCdt1 deficient (4 hours with 500μM IAA) parasites were co-stained with α-Centrin1, α-TgIMC1 and DAPI. A hundred random vacuoles of the parasites were evaluated in three independent experiments. The mean values are plotted on the graphs. The t-test values are listed in TableS4. (C) Immunofluorescent microscopy analysis of RH*ΔKu80TIR1* tachyzoites expressing (-IAA) or TgCdt1^AID-HA^ deficient (+IAA, 4 hours). To determine the number of centrosomes per internal bud, parasites were co - stained with α-Centrin1 (α-mouse IgG Fluor 488), α-TgIMC1 (α-rabbit IgG Fluor 568) antibodies and DAPI (blue). Representative images of the budding parasites and are shown. (D) Quantification of the centrosomes per internal bud in the budding populations of RH*ΔKu80TIR1* tachyzoites expressing (-IAA) or TgCdt1^AID-HA^ deficient (+IAA, 4 hours) based on IFA analysis in panel C. A hundred random vacuoles of the parasites were evaluated in three independent experiments. The mean values are plotted on the graphs. The t-test values are listed in TableS4. (E) Schematics of the *T. gondii* perinuclear structures showing relative position and composition of the bipartite centrosome, centrocone and kinetochores. The drawing depicts one half of the mitotic figure. The dotted line indicates nuclear/cytoplasmic interface. dBC – daughter basal complex. (F) The ultra-expansion microscopy analysis of RH*ΔKu80TIR1* tachyzoites co-expressing TgCdt1^AID-HA^ and TgCEP250-L1^myc^. To visualize the outer core of the centrosome parasites were stained with α-Centrin1 (α-mouse IgG Fluor 568). To detect TgCEP250-L1 in the inner cores of the bipartite centrosome, tachyzoites were stained with α-myc (α-rabbit IgG Flour 488) antibodies. The non-treated tachyzoites (-IAA) retain tightly linked centrioles in mitosis, while the TgCdt1 deficiency (+IAA, 4 hours) led to centrioles disengagement and reduplication. (G and I) Western Blot analysis of the TgCEP250-L1^myc^ expression in the TgCdt1 expressing (-IAA) and deficient (+IAA, 4h) parasites. Western blots were probed with α-myc to identify TgCEP250-L1, α-HA to verify TgCdt1 downregulation and α-TubA to confirm equal loading of the total lysates. The panel I is an overexposed image of TgCEP250-L1^myc^ that shows accumulation of the degradation products. The densitometry analysis of the image is shown on the right (ImageJ). (H) The ultra-expansion microscopy images of RH*ΔKu80TIR1* tachyzoites co-expressing TgCdt1^AID-HA^ and TgCEP250-L1^myc^. The inner core of the centrosome was detected with α-myc (α-rabbit IgG Flour 568) antibodies, the parasite surface with α-IMC1 (α-mouse IgG Fluor 488) and nucleus with DAPI stain. The inner core changes caused by TgCdt1 depletion (+IAA, 4 hours) are indicated. (J) Western blot analysis of RH*ΔKu80TIR1* TgCdt1^myc^ protein extracts non-treated and treated with FAST-AP alkaline phosphatase. Western blots were probed with α-myc to identify TgCdt1 and α-GRA7 to confirm equal loading of the total lysates. Non-phosphorylated (blue arrow) and hyper-phosphorylated (red arrow) TgCdt1 are indicated. (K) IFA analysis of TgCdt1^myc^ localization in RH*ΔKu80TIR1* parasites expressing (-IAA) or deficient for TgCrk4^AID-HA^ (+IAA, 4 hours). Nuclear localization of TgCdt1^myc^ was detected with α-myc (α-rabbit IgG Fluor 568) antibodies and DAPI stain (blue). (L) Western blot analysis of TgCdt1^myc^ in RH*ΔKu80TIR1* parasites expressing (-IAA) or deficient for TgCrk4^AID-HA^ (+IAA, 4 hours). Western blots were probed with α-HA antibodies to confirm TgCrk4 downregulation, α-myc antibodies to show accumulation of the hyperphosphorylated TgCdt1 in TgCrk4-deficient parasites (+IAA), and α-GRA7 to confirm equal loading of the total lysates. Non-phosphorylated (blue arrow) and hyper-phosphorylated (red arrow) TgCdt1 are indicated.

Further examination revealed that TgCdt1 retained the conventional Cdt1 role in control of chromosome rereplication and segregation, and an unexpected role in regulating centrosome reduplication. Echoing the DNA rereplication defect, over 80% of budding TgCdt1-deficient parasites contained doubled number of the Centrin1-positive structures per internal bud (Fig. 7C and D). The ultra-expansion microscopy analysis identified these structures as disengaged centrioles, suggesting premature disengagement. In conventional eukaryotes, two centrioles, constituting the centrosome core, remain joined throughout most of the cell cycle and only briefly disengage in the G_1_ phase, which allows templating and growth of the daughter centriole (centriole duplication) [49]. Similarly, the *T. gondii* centrioles located in the outer core of the bipartite centrosome remain tightly linked during S-phase and mitosis (Fig. 7E and F, -IAA panel, red). However, centrioles in the TgCdt1-deficient tachyzoites at the similar cell cycle stage had drifted a distance from each other (Fig. 7F, +IAA panel, red). In some cases, the separation induced early duplication of the centrioles in mitotic parasites (Fig. 7F, +IAA panel, reduplicated centrioles). Previous studies showed that the outer centrosomal core promotes the assembly of internal buds [16]. In a good agreement with the prediction, tachyzoites lacking TgCdt1 formed multiple internal buds, suggesting that individual centrioles are equally efficient in supporting daughter scaffold assembly (Fig. S5A).

To find out how the TgCdt1 deficiency affected the inner core of the bipartite *T. gondii* centrosome, we endogenously tagged the compartment marker TgCEP250-L1^myc^ in the TgCdt1 AID line (Fig. 7E and F, green channel). Examination of the TgCdt1 depleted parasites revealed dramatic effect on the inner core of the centrosome. While normally dividing the S-phase and mitotic tachyzoites sequentially duplicated and segregated inner cores, the TgCdt1 deficient tachyzoites increased the number of the TgCEP250-L1 positive structures (Fig. 7F, green; Fig. 7H, red; Fig. S5B, red). To confirm the inner core amplification, we compared TgCEP250-L1 levels in parasites expressing or deficient in TgCdt1 (Fig. 7G and I). Contrary to our predictions, we detected the substantial reduction of the full-length TgCEP250-L1 in the IAA treated populations. Interestingly, the reduction of the full-length protein was accompanied by accumulation of the TgCEP250-L1 degradation products, suggesting that amplified TgCEP250-L1-positive structures in TgCdt1 deficient tachyzoites contained truncated TgCEP250-L1 (Fig. 7I, intensity plots). We also examined whether TgCdt1 knockdown affected nuclear compartments linked to the bipartite centrosome (Fig. S5C). To do so, we engineered a TgCdt1 AID line expressed inner kinetochore marker Tg56^myc^ and found that tachyzoites lacking TgCdt1 displayed normal dynamics of kinetochores segregation coordinated with centrocone expansion and resolution (TgMORN1 marker) (Fig. S5D).

Altogether, our analysis of mitotic structures in the TgCdt1 deficient parasites revealed a pronounced effect of the TgCdt1 expression on the outer and inner cores of bipartite centrosome. We determined that TgCdt1 was required for tight connection of the centrioles pair after initial duplication in the G_1_ phase. The observed TgCdt1-dependent proteolysis of TgCEP250-L1 implied that TgCdt1 also plays role in stabilization of the inner core of the centrosome. Together with the DNA replication defects, our results suggested that TgCdt1 controls the “copy once per cell cycle” rule of the chromosome and centrosome duplication in *T. gondii*.

### TgCdt1 is regulated by cell cycle dependent phosphorylation

Evaluation of TgCdt1 expression by Western Blot analysis revealed that, independent of fused epitope, TgCdt1 migrated as two prominent bands (Fig. 6E and Fig. 7J). The apparent molecular weight of the faster moving species corresponded to the predicted TgCdt1 molecular weight of 135kDa, suggesting that the slower moving band represented TgCdt1 containing post-translational modifications. Since phosphorylation affects protein mobility and Cdt1 is known to be regulated by different kinases, we predicted that the slower moving TgCdt1 species was a phosphorylated form of the protein [47, 50]. To test our prediction, we treated the total protein extracts with alkaline phosphatase. Confirming our predication, phosphatase treatment did not affect the lower molecular weight band, but eliminated the upper TgCdt1 band, which validated the existence of a hyper-phosphorylated form of TgCdt1 (Fig. 7J).

Our proteomics studies detected TgCdt1 as a dominant interactor of the G_2_ complex TgCrk4-TgCyc4. To find out if TgCrk4 had any effect on the TgCdt1 expression or localization, we introduced TgCdt1^myc^ into TgCrk4 AID transgenic parasites. A TgCrk4-dependent G_2_ block did not significantly affect TgCdt1 expression or localization (Fig. 7K). Metazoans use a combination of hyper-phosphorylation and geminin inhibition to strategically block Cdt1-dependent DNA replication licensing during G_2_ phase [47]. To find out if TgCdt1 phosphorylation state changes in parasites arrested in G_2_, we analyzed the state of TgCdt1 in both asynchronously dividing and G_2_-arrested parasites (Fig. 7L). G_2_ arrest induced by TgCrk4 depletion led to accumulation of hyper-phosphorylated TgCdt1, confirming cell cycle-dependent regulation of TgCdt1 by post-translational modification. Since hyper-phosphorylated TgCdt1 accumulated in TgCrk4-deficient parasites, it is likely that kinases other than TgCrk4 contribute to TgCdt1 hyper-phosphorylation.

## DISCUSSION

In the current study, we made several critical discoveries regarding the apicomplexan cell cycle. First, we identified the previously unrecognized G_2_ period which operates during apicomplexan endodyogeny. Second, we showed that apicomplexan G_2_ phase is regulated by the Crk4-Cyc4 complex that exists only in parasites dividing in a binary fashion. Third, we identified a *T. gondii* ortholog of DNA replication licensing factor Cdt1 and showed that TgCdt1 is a major interactor of the TgCrk4-TgCyc4 complex. Furthermore, we presented evidence that TgCdt1 has a significantly diminished role in the conservative licensing of DNA replication in G_1_ phase, and primarily controls DNA rereplication and centrosome reduplication in G_2_ phase. Our findings support the cooperative action between TgCrk4-TgCyc4 complex and TgCdt1 to repress multinuclear division, which is likely the default cell cycle mechanism in Apicomplexa phylum. Thus, contrary to the common view of *T. gondii* endodyogeny as the simplest mode of apicomplexan division, our findings strongly argue that endodyogeny has more complex regulation of the cell cycle and had instead evolved additional mechanisms to suppress DNA rereplication cycles.

Although rare, the multinuclear division is not a novelty in eukaryotes. The best-known example of multinuclear division in higher eukaryotes is in the embryonic development of *Drosophila melanogaster* [51]. The first thirteen cycles of DNA amplification in the fruit fly zygote consists of S-phase (chromosome replication) and mitosis (chromosome segregation) repeating to produce a single cell with 500 nuclei, which is then called a blastoderm. Binary division is then introduced with the G_2_ phase, which extends the cell cycle and allows the time for the first cytokinetic event to occur and separate these nuclei by cell membranes. Thus, one of the critical functions of G_2_ is to repress multinuclear division by enforcing the “once per cell cycle duplication” rule that minimizes the chances of losing or altering genetic information due to chromosome mis-segregation. *Drosophila* development is somewhat reminiscent of cell division in apicomplexan parasites. Given the fact that only a fraction of apicomplexans divides by a binary mode, and that the vast majority undergo multinuclear replication, it is tempting to suggest that multinuclear division is the ancestral mechanism of apicomplexan cell division, with binary division evolving later to accommodate the specific needs of such apicomplexans genera as *Toxoplasma*, *Hammondia*, *Neospora*, *Besnoitia*, and *Babesia* [3]. In support of this proposition, reflecting the additional levels of cell cycle regulation, *T. gondii* maintains an extended repertoire of cell cycle machinery to predominantly divide into two [2, 6]. What mechanism would apicomplexan parasites evolve to convert from multinuclear to binary division? The zygotic fruit fly example points toward the possibility that some apicomplexans evolved a mimic G_2_ period by installing mechanisms to repress relicensing of DNA replication and centrosome duplication and to advance cytokinesis (budding). This could be achieved by connecting replicated chromosomes to the cytoskeleton of future daughter buds using a modular centrosome such as the bipartite *T. gondii* centrosome [16]. The inner centrosomal core is functionally linked to chromosome segregation and the outer core is involved in budding, and, during endodyogeny, they are permanently associated with one another [15, 16, 19]. It has been shown that separating centrosomal cores leads to mitotic death in tachyzoites [15, 16].

Characterizing the processes regulated by TgCrk4 led to the construction of the G_2_ network in *T. gondii*, which includes its immediate effectors, direct interactors, and cellular pathways that uniquely respond to the G_2_/M transition (Fig. 8). We identified the families of conventional G_2_ regulators, such as components of the origin recognition complex (TgORC4), the anaphase-promoting complex APC/C (Cdc20), and a DNA replication licensing factor (TgCdt1). However, the immediate effectors differed from known G_2_ regulators, and often included parasite-specific groups of proteins. For example, gene expression seems to be directly regulated by the phosphorylation of a parasite-specific AP2X-9 DNA binding factor. Control of centrosome reduplication involves the direct phosphorylation of the centrosomal protein TgCEP530, Ca^++^ dependent kinase TgCDPK7, as well as interaction with a highly deviated Cdt1 ortholog [20, 52]. Although their direct effectors are not yet known, progression through the G_2_/M checkpoint has a significant effect on the IMC components of daughter buds (TgIMC16, TgIMC30, TgIMC33, TgIMC34, TgIMC36), Apicomplexa-specific components of kinetochores (TgAKiT1, TgAKiT6), and the bipartite centrosome (TgCEP250-L1, TgCEP250, TgCEP530) [15, 16, 19, 20, 27–29, 36]. Importantly, this novel G_2_ network incorporates factors that were previously shown to induce reduplication of centrosomes and nuclear DNA. Among them are the deubiquitinase TgOTUD3A and centrosomal kinase TgMAPK1 [16, 31].

**Figure 8.**
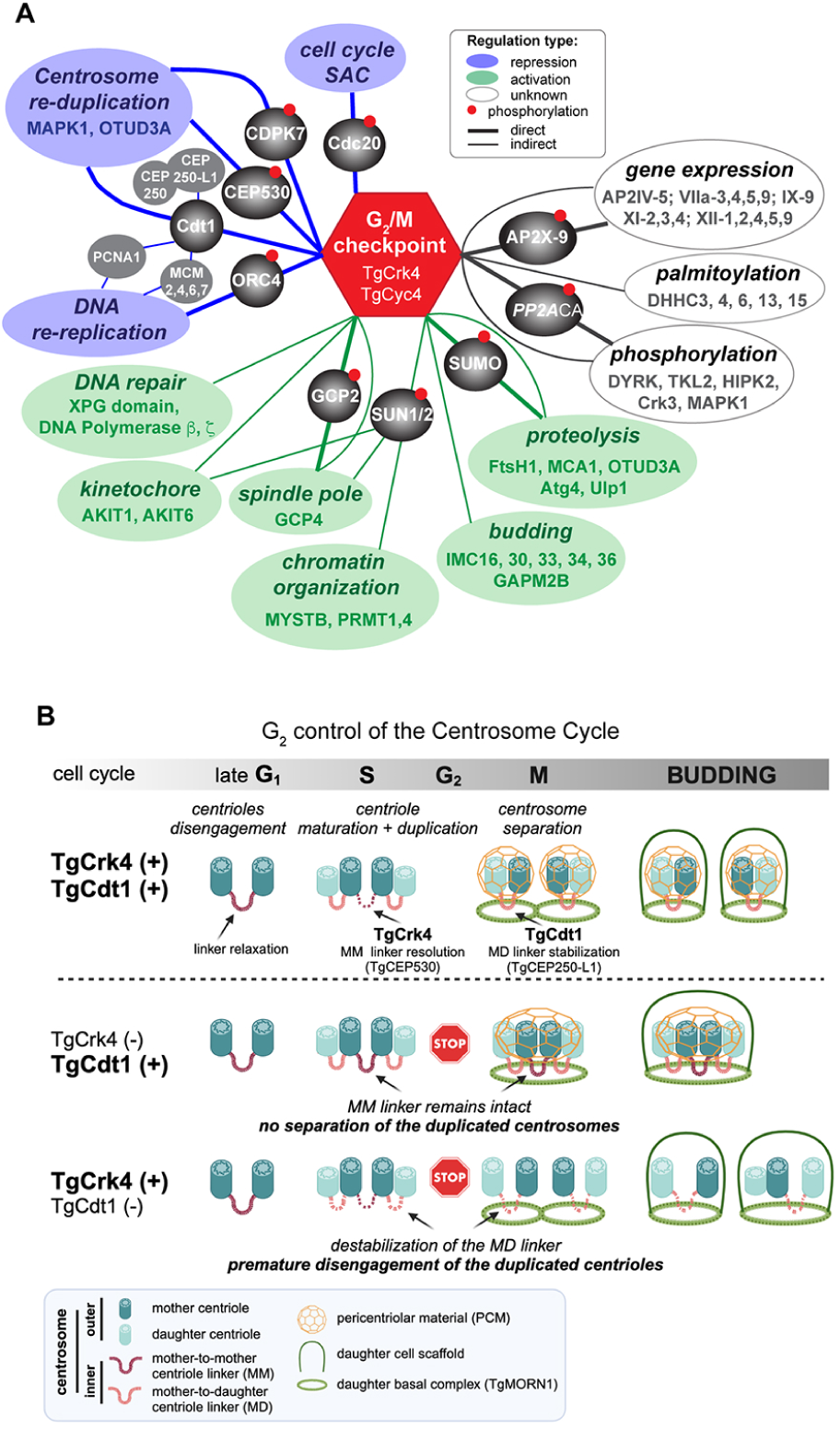
The mechanisms that regulate G_2_ phase in *T. gondii* endodyogeny. (A) The summary of the *Toxoplasma* G_2_/M network identified in this study. The novel TgCrk4-TgCyc4 complex regulates *T. gondii* G_2_/M checkpoint, shown as the red STOP sign. The affected cellular processes are indicated by ovals. TgCrk4-dependent phosphorylation of TgCDPK7, TgORC4 and TgCEP530 likely represses centrosome reduplication and chromosome rereplication (blue ovals), which also includes dominant interaction with DNA replication licensing factor TgCdt1. Activation of the spindle pole assembly and targeted proteolysis is mediated by phosphorylation of TgGCP2 and TgSUMO protease, respectively (green ovals). The G_2_/M transition also controls APC/C activation via direct phosphorylation of TgCdc20 (blue oval), while regulators of the kinetochore assembly and concurrent assembly of internal buds are currently unknown (green ovals). The G_2_ phase has a major effect on global protein phosphorylation and gene expression, likely via direct phosphorylation of TgPP2ACA and TgAP2X-9. (B) Putative mechanism of regulation of the *Toxoplasma* centrosome cycle. The upper panel illustrates the normal centrosome dynamics during tachyzoite cell cycle, which progression is indicated on the top grey bar. The cartoon shows two cores of the bipartite centrosome and the components of the daughter cell scaffold. The outer core of the centrosome contains two parallel centrioles. We propose that the inner core structurally resembles a proteinaceous fiber that connects centrioles in the outer core and is made of coil-coiled domain containing proteins of the CEP family. Major steps of the centrosome cycle include centriole disengagement in G_1_, duplication and maturation in S-phase, and centrosome separation at G_2_/M transition. The changes to the outer and inner core of the bipartite centrosome caused by TgCrk4 and TgCdt1 deficiency are shown on the two bottom panels. The STOP sign indicates a position of the TgCrk4 regulated G_2_/M checkpoint.

One of the unexpected findings from our studies is the dominant interaction of TgCrk4-TgCyc4 complex with an ortholog of the DNA replication licensing factor, TgCdt1. Eukaryotic Cdt1 is a multifunctional protein that plays distinctive roles in nearly all cell cycle phases [42]. Cdt1 activates pre-replication complexes in G_1_ phase, is actively removed from the S-phase nucleus, and kept off the chromatin in G_2_ phase to avoid chromosome rereplication. Cdt1 also plays a role in kinetochore assembly by securing the connection of Ndc80 to spindle fibers in mitosis, thereby assisting in chromosome segregation [46]. Our study found similar features as well as substantial differences between *T. gondii* Cdt1 and its eukaryotic counterpart, which share limited sequence similarity. First, TgCdt1 was detected in the nucleus of S, G_2_, and mitotic parasites, which is opposite to temporal expression of Cdt1 in mammalian cells and yeast. Second, as a dominant partner of the TgCrk4-TgCyc4 complex, TgCdt1 played its primary roles in G_2_ and mitosis. In this aspect, the TgCrk4-TgCyc4 complex operates in an analogous manner to the mammalian Cdk2-Cyclin A complex. However, the Cdk2-Cyclin A complex is dispensable for somatic cell division, while TgCrk4 and TgCyc4 are independently required for tachyzoite survival [41]. The functional resemblance of TgCrk4-TgCyc4 and Cdk2-Cyclin A complexes was also supported by the structural similarities between the cyclin partners. Third, we found that depleting TgCdt1 had no sizable effect on licensing of DNA replication in G_1_ phase, suggesting alternative mechanisms of regulation of DNA replication in parasites. The fact that TgCdt1 is absent during primetime of the Cdt1 function in conventional eukaryotes (late mitosis and half of G_1_ phase) further supports the minimal involvement of TgCdt1 in licensing DNA replication in G_1_ phase. Furthermore, contradicting its expected role as a licensing factor, TgCdt1 was constitutively present in the nucleus while DNA replication was ongoing (S-phase).

How does the TgCdt1 factor contribute to the G_2_ phase progression? Based on our collective findings we proposed a model of the centrosome cycle regulation by TgCdt1 and G_2_ kinase TgCrk4 during *T. gondii* endodyogeny. The centrosome cycle consists of sequentially executed steps of the centrioles disengagement (G_1_), centriole duplication and maturation (S-phase), and separation of the duplicated centrosomes each containing a pair of centrioles (G_2_ and mitosis) (Fig. 8B). The cell cycle dependent assembly and resolution of the linker controls centrioles connection [49]. In the conventional cell cycle, a single linker connecting centrioles is assembled in G_1_ and is resolved in G_2_ phase to allow centrosome separation. The orthogonal position of the daughter centriole likely inhibits early disengagement from the mother centriole in mitosis. However, *Toxoplasma* centrioles lay parallel and may require additional connectors to hold mother and newly assembled daughter centriole during S, G_2_ phases and mitosis, thus, preventing an unlawful centrioles reduplication [8]. According to our model, unlike a conventional centrosome, the *Toxoplasma* centrosome uses two connectors that differentiate a mother-mother and a mother-daughter pair of centrioles (Fig. 8B). One linker is inherited from the previous cycle. It connects two mother centrioles in G_1_, S and G_2_ phases. The other linker, formed in S-phase, holds mother and daughter centrioles until the cell division is complete. The inheritance of the centrioles from the previous to the current cycle suggests that the mother-daughter linker maturates into a mother-mother linker as the parasite enters the new cell cycle. Judging by the joined centrosome phenotype, TgCrk4 likely regulates resolution of the mother-mother centriole linker, possibly by direct phosphorylation of TgCEP530 located in the uncharacterized intermediate centrosomal core [20]. Previous studies implicated mitotic kinase TgNek1/2 in the process of centrosome separation [53]. Since we could not detect TgNek1/2 in the G_2_ network, this kinase likely acts upstream of TgCrk4. Contrary to TgCrk4 knockdown phenotype, TgCdt1 deficient tachyzoites prematurely disengaged centrioles in G_2_ and mitotic parasites, suggesting that TgCdt1 regulates the mother-daughter centriole linker. The likely mechanism involves stabilization of another coil-coiled domain protein, TgCEP250-L1 [16, 19]. Our model implies that differential protein composition of the mother-mother and mother-daughter centriole linkers facilitates cell cycle-dependent processing of individual connectors. For instance, the presence of TgCEP530 in the mother-mother and not mother-daughter connector can make the mother-mother linker accessible for the TgCrk4-dependent resolution in the G_2_ phase. It drives the centrosome separation but preserves the connection of the centrioles within individual centrosomes. On the other hand, the alternative TgCEP250-L1 state in the connectors may explain selective protection of the mother-daughter and not mother-mother linker by TgCdt1 in G_2_ phase. Supporting our model, it has been recently shown that in mitotic parasites, the C-Nap related protein TgCEP250 plays a role in connecting the outer core (centrioles) and inner core (linker) [15]. Moreover, similar to TgCEP250-L1, TgCEP250 undergoes proteolytic maturation suggesting co-expression of the factors in a centriole linker. Further studies are required to dissect the mechanisms connecting parallel *T. gondii* centrioles. Comprehensive analysis of the *T. gondii* centrosome cycle will answer outstanding question of the apicomplexan biology such as why only a small group of apicomplexans preserved centrioles, why the parallel orientation is favored, and whether the centriole retention benefits the binary type of cell division.

Lastly, it is tempting to suggest that the unique expression profile of TgCdt1 had evolved to accommodate a special feature of *T. gondii* tachyzoites, which is the ability to egress at any time of intravacuolar division [54]. Apicomplexan parasites are obligatory intracellular organisms that cannot replicate outside of a host cell. Thus, degradation of essential cell cycle machinery such as TgCdt1 may serve as a precautionary measure that potentially egressing parasites do not replicate in an unsafe environment that may not support their growth. A similar removal of DNA licensing factors was reported in *Plasmodium falciparum*, suggesting this is a mechanism common across the phylum [55]. We also showed that TgCdt1 retained a feature that was recently discovered in the mammalian Cdt1: stage-specific hyper-phosphorylation to repress Cdt1-dependent licensing of DNA replication in S and G_2_ phases [47]. This finding together with the discovery of the primary role that TgCdt1 plays in relicensing chromosomes and re-duplicating centrosome raises an important question of how tachyzoites segregate the functions of initiation of DNA replication and relicensing, which can be addressed in future studies.

## MATERIALS AND METHODS

### Parasite Cell Culture and Plaque Assay

*T. gondii* RH *ΔKU80Δhxgprt AtTIR1* and RH *TatiΔKU80* strains were grown in Human Foreskin Fibroblasts (HFF) (ATCC, SCRC-1041) in DMEM media (Millipore Sigma) supplemented with 10% FBS [56, 57]. Cultures were tested for mycoplasma contamination using Mycoplasma Detection kit (MP Biomedicals). The viability of transgenic strains was measured by plaque assays in which monolayers of HFF cells were infected with 50 parasites per well. Cultures were treated with either a 95% EtOH (control), or 500µM Indole-3-acetic-acid (IAA or auxin), or 1µg/mL of anhydrotetracycline (ATc) to trigger protein degradation or transcriptional down regulation, respectively. Plaques developed after 7 days growth at 37°C were fixed, stained, and counted. Three biological replicates of each assay were performed.

### Phylogenetic Analysis

Protein sequences were downloaded from UniProt (www.uniprot.org) database and analyzed using NGPhylogeny (www.NGPhylogeny.fr.) custom workflow [58, 59]. Proteins were aligned with MUSCLE algorithm, curated with Block Mapping and Gathering with Entropy (BMGE) [60, 61]. Maximum likelihood-based inference with Smart Model Selection was implemented using PhyML+SMS algorithm [62]. We applied 100 bootstraps to test optimization of the tree. Phylogenetic trees were constructed using Newick utilities [63]. Amino acid sequences of the analyzed proteins are listed in Table S2.

### Construction of Transgenic Strains

All transgenic strains created in this study and used primers are listed in Table S1. Targeting constructs were verified by PCR using gene-and epitope tag-specific primers, ensuring that tags were incorporated at the appropriate genomic locus.

#### Endogenous C-terminal Tagging and Conditional Expression

To create conditional expression models for TgCrk4 and TgCdt1, we amplified a fragment of the 3’ end of the gene of interest via PCR and cloned it into the pLIC-mAID-3xHA-hxgprt, and/or pLIC-10xHA-CmR, vectors digested with PacI endonuclease using Gibson Assembly method. 50µg DNA of the resulting constructs were linearized with endonuclease within the gene fragment and transfected using Amaxa Nucleofector II device (Lonza) into RH *TIRΔKU80Δhxgprt* and RH *ΔKU80Δhxgprt* parents respectively with 100 µl Cytomix buffer supplemented with 20mM ATP and 50mM reduced glutathione. The same approach was applied in building myc-epitope tagged TgCrk4, TgCdt1 and TgCEP250-L1. Genomic fragments were amplified then cloned into pLIC-3xMyc-DHFR and/or pLIC-10x-MYC-CmR, linearized and transfected into respective parental strains. Dual transgenic strains were developed in the same fashion and sequentially electroporated into parental transgenic strains and selected for with alternative drugs. All transgenic parasites were incubated for 24 hours in regular growth media prior to drug selection, limited dilutions were used to obtain individual clones and incorporation was tested using PCR and IFA analysis.

#### Endogenous N-terminal Tagging and Conditional Expression

To build tet-OFF mutant TgCyc4 a fragment of the 5’ end of the gene was amplified (Table S1 lists primers used) and digested with BglII/NotI for compatible ends and ligated into the promoter replacement vector pTetO7sag4-3xHA-DHFR [6]. Constructs were linearized with endonuclease within the gene fragment and electroporated into RH*TatiΔKu80* strain [57].

### Immunofluorescence Microscopy Analysis

Monolayers of HFF cells were grown on coverslips and infected with parasites under indicated conditions. Cells were fixed with 4% paraformaldehyde, permeabilized with 0.5% Triton TX-100, blocked with a 3% bovine serum albumin solution, and incubated consecutively with primary and secondary antibodies. Primary antibodies used: rat α-HA (3F10, Roche Applied Sciences), mouse α-Centrin1 (clone 20H5; Millipore Sigma), mouse α-Atrx1 (kindly provided by Dr. Peter Bradley, UCLA, CA), rabbit α-Myc (clone 71D10, Cell Signaling), rabbit α-MORN1 (kindly provided by Dr. Marc-Jan Gubbels, Boston College, MA), mouse and rabbit α-IMC1 (kindly provided by Dr. Gary Ward, University of Vermont, VA), mouse α-Tubulin A (kindly provided by Dr. Jacek Gaertig, University of Georgia, Athens GA), and mouse α-Tubulin A (clone 12G10; DSHB). Alexa conjugated secondary antibodies of different emission wavelengths (Thermo Fisher) were used at 1:500 dilution, and nucleic acids were stained with 4’,6-diamidino-2-phenylindole (DAPI, Sigma). Stained parasites on the coverslips were mounted using Immu-mount (Fisher 9990402), dried overnight at 4°C, and viewed on a Zeiss Axiovert Observer Microscope equipped with a 100X objective and ApoTome slicer. Collected images were processed in Zen2.2 and Adobe Photoshop 2020 using linear adjustment when needed.

### Ultra-Expansion Microscopy

The tachyzoite internal structures were examined by ultra-expansion previously described [18]. Briefly, tachyzoites growing in HFF cells were fixed in with 4% PFA, incubated in 2% formaldehyde, 1.4% acrylamide (AA) in PBS for 5 hours at 37 °C. The sample was converted into a gel by incubation in 19% (w/w) sodium acrylate, 10% (w/w) AA and 0.1% (w/w) BIS-AA in PBS supplemented with ammonium persulfate and TEMED (tetramethylethylenediamine) for 1 hour at 37 °C, followed by incubation in denaturation buffer (200 mM SDS, 200 mM NaCl, 50 mM Tris, pH 9) at 95 °C for 90 min. Gels were expanded in ddH_2_O overnight and washed twice in PBS on the next day. Primary and secondary antibody incubation was performed for 3 hours at 37°C. The primary antibodies were used at the following dilution: mouse α-Tubulin A (12G10, DSHB; 1:250), mouse α-myc (9B11, Cell Signaling; 1:500), rabbit α-myc (71D10, Cell Signaling 1:250), rabbit α-IMC1 (kindly provided by Dr. Gary Ward, University of Vermont, VA; 1:250), rabbit α-MORN1 (kindly provided by Dr. Marc-Jan Gubbels, Boston College, MA; 1:100) and mouse α-Centrin1 (clone 20H5; Millipore Sigma; 1:100). Stained gels were washed three times with PBST (1xPBS + 0.1% Tween20) before a second overnight expansion in ddH_2_O.

### Western Blot Analysis

Infected HFF monolayers were lysed in Laemmli sample buffer, sonicated, and heated at 95°C for 10 minutes. Alternatively, lysates were prepared from purified parasites. After separation on SDS-PAGE gels, proteins were transferred onto nitrocellulose membranes and probed with rat α-HA (3F10, Roche), rabbit α-Myc (Cell Signaling), rabbit α-TgPCNA2 (kindly provided by Dr. Michael White, UC Riverside, CA), mouse α-GRA7 (kindly provided by Dr. Peter Bradley, UCLA CA) and mouse α-Tubulin A (12G10; kindly provided by Dr. Jacek Gaertig, University of Georgia, Athens GA). After incubation with a-species specific secondary horseradish peroxidase (HRP)-conjugated antibodies (Jackson ImmunoResearch), proteins were visualized by enhanced chemiluminescence detection (MilliporeSigma Immobilon). For western blot analysis of TgCdt1 phosphorylation parasites were collected, washed in PBS with protease and Halt^TM^ phosphatase inhibitor cocktail, resuspended in PBS and treated with FAST AP according to manufacturer recommendation (NEB).

### FACS Scan Analysis

Parasites DNA content was evaluated by flow cytometry using propidium iodide staining of tachyzoites DNA. HFF Monolayers were infected at 1:1 MOI and left to grow overnight at 37°C. Next day, TgCrk4AID-HA parasites were treated with 500µM Auxin for 0, 4 hr, with 1 hour and two-hour recovery periods after auxin was removed, and TgCdt1AID-HA parasites were treated with 0, 4hr, 6hr, and 8hr of 500µM Auxin. Parasites were collected by scraping, passing through an 18G needle 5X, then filtering with a 3µm filter. Parasites were pelleted by centrifugation at 1800rcf for 15min. Media was aspirated, and parasites were resuspended in 300µL of cold PBS with 700µL of ice cold 100% molecular grade ethanol added while being vortexed. Samples were then stored at -20°C until day of FACS scan analysis. Day of, parasites were pelleted at 3000rcf for 20min, ethanol was aspirated, and pellet was resuspended in 900mL of 50µM Tris solution pH 7.5. 100µL of 2mg/mL propidium iodide (Fisher) for a final concentration of 0.2mg/mL, and 20µL of 10mg/mL RNase A for a total concentration of 0.2mg/mL. Samples were left to incubate in the dark at room temperature for 30 minutes. DNA content was measured based on the intensity of the emission from PI-stained DNA using the PE-Texas Red-A laser. Parameters for the specific size of Toxoplasma was used to determine single cells from debris and non-single cells, percentages of each cell cycle phase were calculated based on the defined gates for each population.

### Proteomics analysis

#### The immune-precipitation sample preparation

To analyze TgCrk4, TgCyc4 and TgCdt1 complexes, the samples were prepared from 10^9^ parental (RH*ΔKu80Δhxgprt AtTIR1* or RH Tati *ΔKu80*) and tachyzoites expressing endogenously tagged TgCrk4^AID-HA^, ^HA^TgCyc4 or TgCdt1^myc^ as previously described [11]. Parasites were collected by filtration and centrifugation. Extracted proteins were incubated with α-HA or α-myc magnetic beads (MblBio). The efficiency of immune precipitation was verified by Western blot analysis. Protein samples were then processed for mass spectrometry-based proteomic analysis as previously described [64, 65].

#### The global proteome analysis sample preparation and mass-spectrometry

Samples for TgCrk4 global proteome were prepared and processed as previously described [11]. Briefly, tachyzoites were collected and solubilized with 5% SDS (w:v) in 50mM TEAB. Equal amounts of protein (200μg) was fractionated and were processed for LC-MS/MS using s-traps (Protifi). Phosphopeptides of each fraction were enriched on the TiO_2_ nanopolymer beads (Tymora Analytical), eluted, dried and resuspended in H_2_O/1% acetonitrile/0.1% formic acid for LC-MS/MS analysis. Peptides were analyzed on a hybrid quadrupole-Orbitrap instrument (Q Exactive Plus, Thermo Fisher Scientific). Full MS survey scans were acquired at 70,000 resolution. The top 10 most abundant ions were selected for MS/MS analysis.

#### Raw data processing

Files were processed in MaxQuant (v 1.6.14.0, www.maxquant.org) and searched against the current Uniprot *Toxoplasma gondii Me49* protein sequences database. Search parameters included constant modification of cysteine by carbamidomethylation and the variable modifications, methionine oxidation, protein N-term acetylation, and phosphorylation of serine, threonine, and tyrosine. Proteins were identified using the filtering criteria of 1% protein and peptide false discovery rate. Protein intensity values were normalized using the MaxQuant LFQ function [66].

Label free quantitation analysis of the global proteome and phosphoproteome was performed using Perseus (v 1.6.14.0), software developed for the analysis of omics data [67]. LFQ Intensity values were Log2-transformed, and then filtered to include proteins containing at least 60% valid values (reported LFQ intensities) in at least one experimental group. Then, the missing values in the filtered dataset were replaced using the imputation function in Perseus with default parameters [67]. Statistical analyses were carried out using the filtered and imputed protein groups file. Statistically significant changes in protein abundance are determined using Welch’s t-test p-values and z-scores.

### Statistical analysis

#### Global protein expression and phosphorylation changes

To determine a relative abundance of the phosphorylated peptides, each phosphorylation intensity value was normalized to intensity of corresponding protein expression (global proteome). We then introduced an improved Gaussian model to identify proteins or phosphorylated peptides that were significantly altered during checkpoint block induced by TgCrk4 degradation (4 hours with auxin) [68–71]. In brief, a gaussian distribution was implemented to model the log2 fold change (log2FC) of normalized protein/peptide intensities. Using a sliding window from low to high of average protein/peptide intensity, we modeled the variance changing of log2FC and selected proteins/peptides within a specific range of intensity. For each replicate, the maximum likelihood estimation (MLE) was used to estimate the parameters in Gaussian distribution from each sliding window after removing entities with log2FC higher than the top 95% or lower than the bottom 5%. The p-value was calculated as the probability of the fitted Gaussian distribution higher or lower than the observed log2FC value when log2FC was higher or lower than the fitted expectation. The false discovery rate (FDR) was used to adjust p-value. We selected significantly changed entities based on the criteria that log2FC > 1 and adjusted p-value < 0.1, or log2FC < -1 and adjusted p-value < 0.1 consistently in all the replicates.

#### Heatmap construction

To capture phosphorylation pattern changes due to TgCrk4 deficiency, we compared pairwise significantly changed phosphorylated peptides detected in untreated, auxin treated for 30mins, or 4 hours parasites. We started with building K-means classification for the 0/30 mins pair. Akaike information criterion (AIC) and BIC were used to search optimized category numbers of K-means classification. The normalized intensity of each selected peptide was further divided by the maximum intensity of this peptide from checkpoint block time to 4 hours, which is represented as ‘Proportion-of-max’ in heatmap. The enriched GO terms for each group of genes are identified by using a hypergeometric test compared with whole *T. gondii* genome with FDR adjusted p-value lower than 0.1.

#### Phosphorylation motif search

The featured motifs of phosphorylated peptides were analyzed by using motif-x algorithm Soft MoMo v5.1.1 (http://meme-suite.org/tools/momo) [72]. For analysis, we selected 13-mer phosphorylated peptides with phosphor modified amino acid residue positioned between 6 residues upstream and downstream of the phosphorylation site. In cases where flanking restudies were missing in the MS identified peptide, the neighboring sequence was extracted from the predicted protein sequence in *T. gondii* genome (ToxoDB). The background or control datasets were selected according to the type of amino acids phosphorylated in query data. The motif sequence was considered when the minimum number of occurrences was over 25 and P value < 1e^-9^.

### Structural analysis

Using UniProt database, amino acid sequences were downloaded and used to build protein homology models in the SWISS-MODEL suite [73]. The resulting models were further analyzed in PyMol (www.pymol.org). Available images of the folded proteins were downloaded from AlphaFold2. Alignment of the protein sequences downloaded from UniProt database was performed using MUSCLE algorithm and exported to Jalview [74].

## SUPPLEMENTAL DATA

**Table S1.** Transgenic strains and primers used in the study.

**Table S2.** Entries used in phylogenetic analysis.

**Table S3.** Proteomics data (IP/MS).

**Table S4.** Raw counts used in the figures.

**Table S5.** MaxQuant analysis of TgCrk4 global proteome and phosphoproteome.

**Table S6.** Analysis of TgCrk4 global proteome.

**Table S7.** Analysis of TgCrk4 global phosphoproteome.

**Table S8.** Search for G_2_-dependent phosphorylation motifs and TgCrk4 substrates.

---

**Figure S1.**
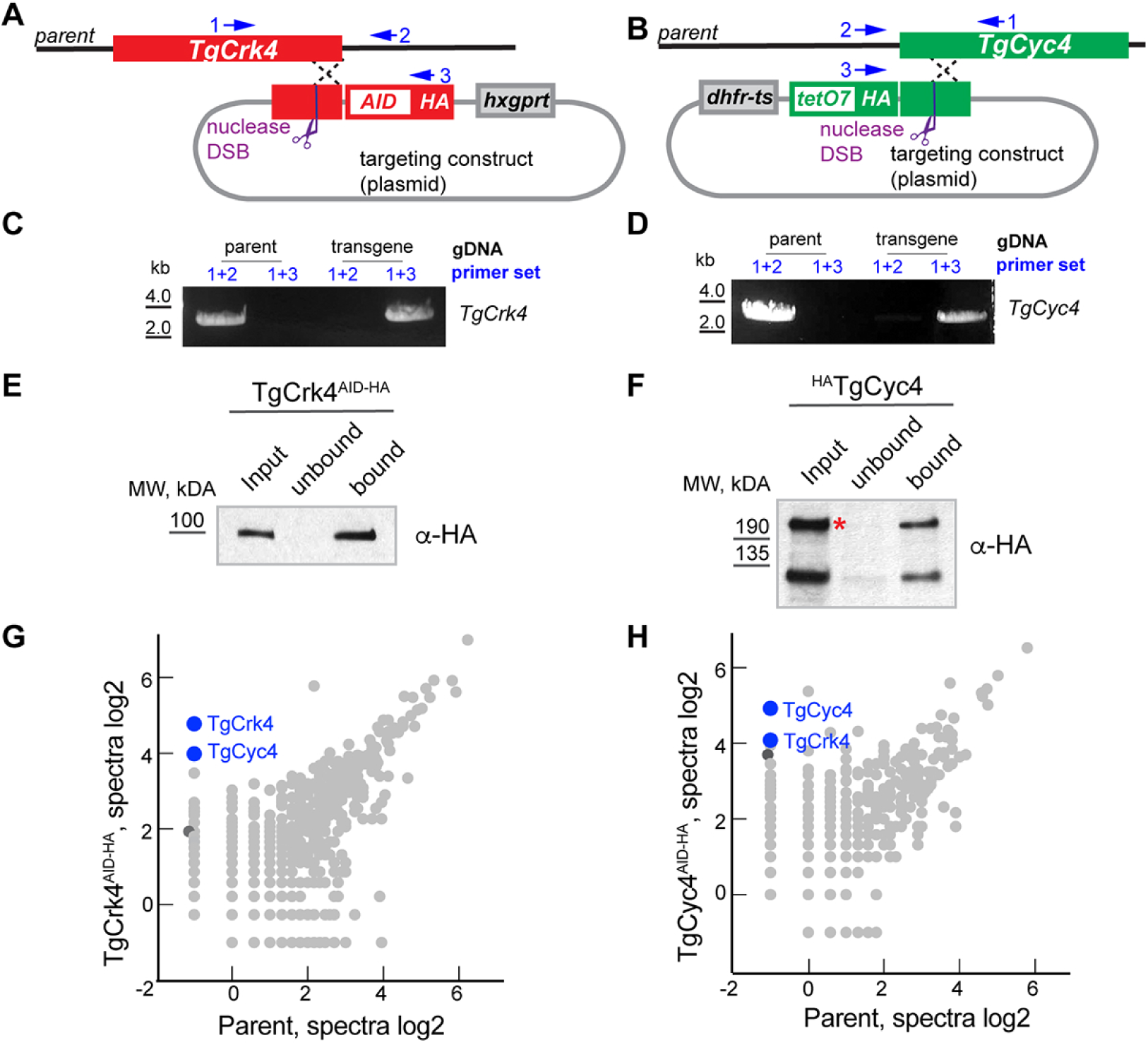
Generation and analysis of conditional expression models for TgCrk4 and TgCyc4. (A) Schematics for constructing TgCrk4 AID-modified gene. Targeting plasmid included 3’ fragment of *TgCrk4* genomic locus fused with encoded sequence for mini-version of AID (mAID), 3xHA (HA) epitopes and the drug-selection marker *hxgprt* gene (grey box). The plasmid linearization with a unique endonuclease induced recombination at the *TgCrk4* locus. Schematics also indicates the relative position of the primers used to conform the TgCrk4 knock-in (panel C). (B) Schematics for constructing TgCyc4 Tet-OFF-modified gene. The 5’ fragment of *TgCyc4* genomic locus was amplified and cloned into the targeted plasmid to create N-terminal fusion with 3xHA (HA) epitopes (grey box). The plasmid linearization with a unique endonuclease induced recombination at the *TgCyc4* locus. Schematics also indicates the relative position of the primers used to conform the TgCyc4 knock-in (panel D). (C and D). PCR analysis of the parental and transgenic lines expressing TgCrk4^AID-HA^ or ^HA^TgCyc4. The combination of the primers used to detect either native or recombined locus are shown. (E and F). Western Blot analysis of TgCrk4 (E) and TgCyc4 (F) immunoprecipitation. The protein complexes were immunoisolated from the soluble fraction [In] (input) of parasites co-expressing endogenous TgCrk4^AID-HA^ or ^HA^TgCyc4. Beads with precipitated complexes [B] and depleted soluble fraction [Un] (unbound) were probed with α-HA (α-rat IgG-HRP) antibodies to confirm efficient pulldown of the target proteins. (G and H) The results of the SAINT analysis of TgCrk4 (G) and TgCyc4 (H) proteomes. The log2 values of the protein spectra detected by mass-spectrometry analysis of the parent parasites and parasites expressing TgCrk4^AID-HA^ or ^HA^TgCyc4 are plotted on the graph. The TgCrk4-TgCyc4 complex is indicated with blue color. TGME49_247040 protein is shown as a dark grey circle.

**Figure S2.**
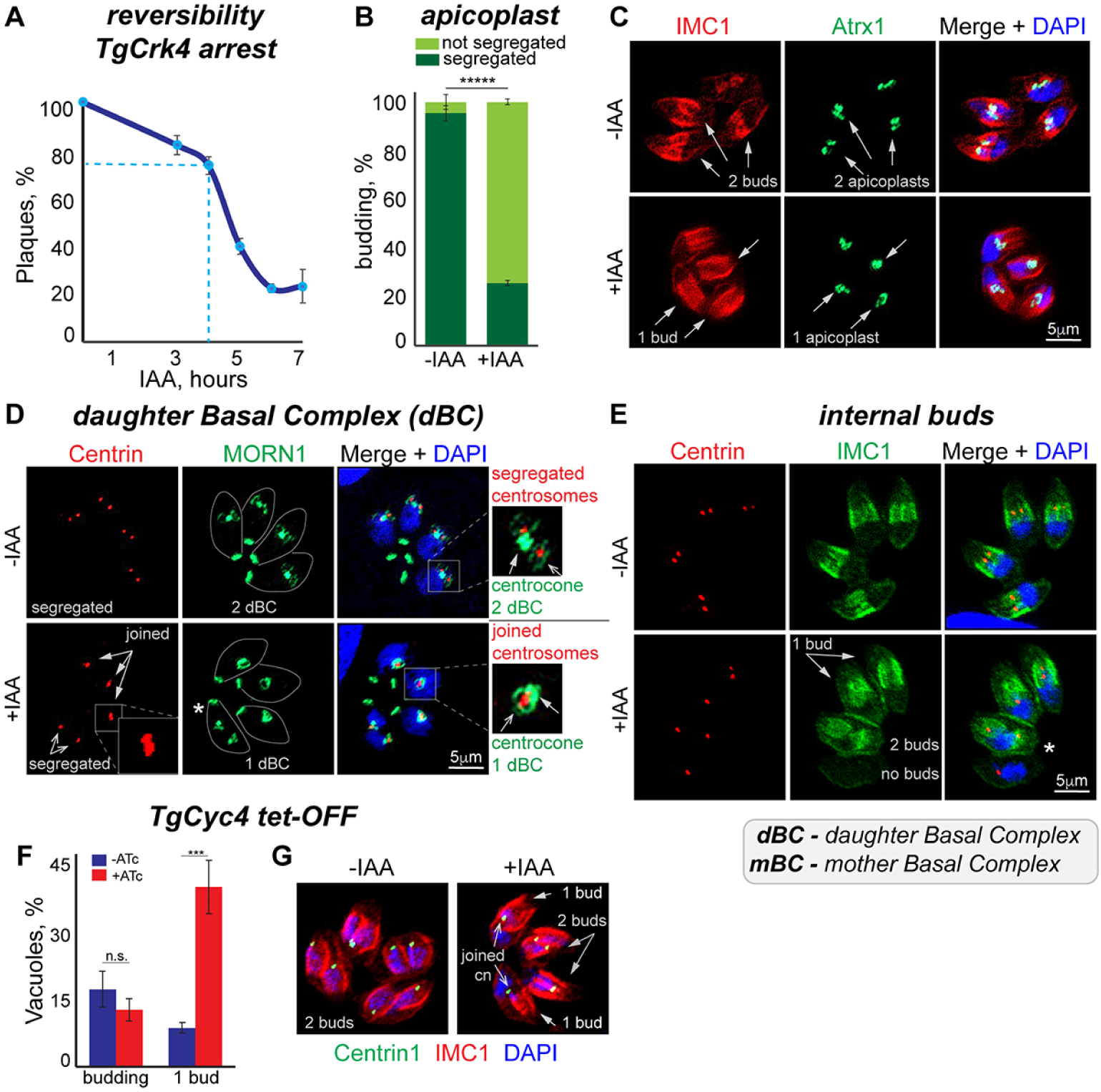
Characterization of TgCrk4 knockdown phenotype. (A) The reversibility of the TgCrk4 induced block was determined by plaque assay. Freshly invaded RH TgCrk4^AID-HA^ parasites were incubated with 500μM IAA for indicated times before the medium was replaced with normal growth medium without IAA to allow for plaque development. The plotted values represent the average plaque numbers from three independent measurements. (B) Quantification of the apicoplast segregation defect caused by RH TgCrk4^AID-HA^ deficiency. The number of the vacuoles containing budding parasites that segregated or did not segregate apicoplast were quantified in non-treated and treated with 500μM IAA for 8 hours parasites using TgAtrx1 staining as shown on panel C. The longer IAA treatment allowed development of the bigger buds to aid quantifications. A hundred vacuoles of the budding parasites were examined in three independent experiments. The mean values are plotted on the graph. The t-test values are listed in TableS4. (C, D and E) Immunofluorescent microscopy analysis of RH*ΔKu80TIR1* TgCrk4^AID-HA^ tachyzoites incubated without (-IAA) or with 500μM IAA for 4 hours (+IAA). (C) Parasites were co-stained with α-TgAtrx1 (α-mouse IgG Fluor 488), α-TgIMC1 (α-rabbit IgG Fluor 488) antibodies and DAPI to visualize apicoplast, internal buds and nuclei. (D) Parasites were co-stained with α-TgMORN1 (α-rabbit IgG Fluor 488), α-Centrin1 (α-mouse IgG Fluor 488) antibodies and DAPI to determine the number of centrosomes and daughter basal complexes (dBC). (E) Parasites were co-stained with α-TgIMC1 (α-rabbit IgG Fluor 488), α-Centrin1 (α-mouse IgG Fluor 488) antibodies and DAPI to visualize internal buds and determine the number of centrosomes per parasite. (F) Quantification of the defects caused by TgCyc4 downregulation. The TgCyc4 Tet-OFF parasites were grown without or with ATc for 16h. The average number of the vacuoles containing budding parasites and parasites forming a single bud in three independent experiments is plotted on the graph. (G) The immunofluorescence microscopy analysis of the TgCyc4 Tet-OFF parasites grown without or with ATc for 16h. The parasites were co-stained with α-Centrin1 (α-mouse IgG Fluor 488), α-TgIMC1 (α-rabbit IgG Fluor 488) antibodies and DAPI to visualize centrosomes, internal buds and nuclei. The quantifications of three independent experiments are shown on panel F.

**Figure S3.**
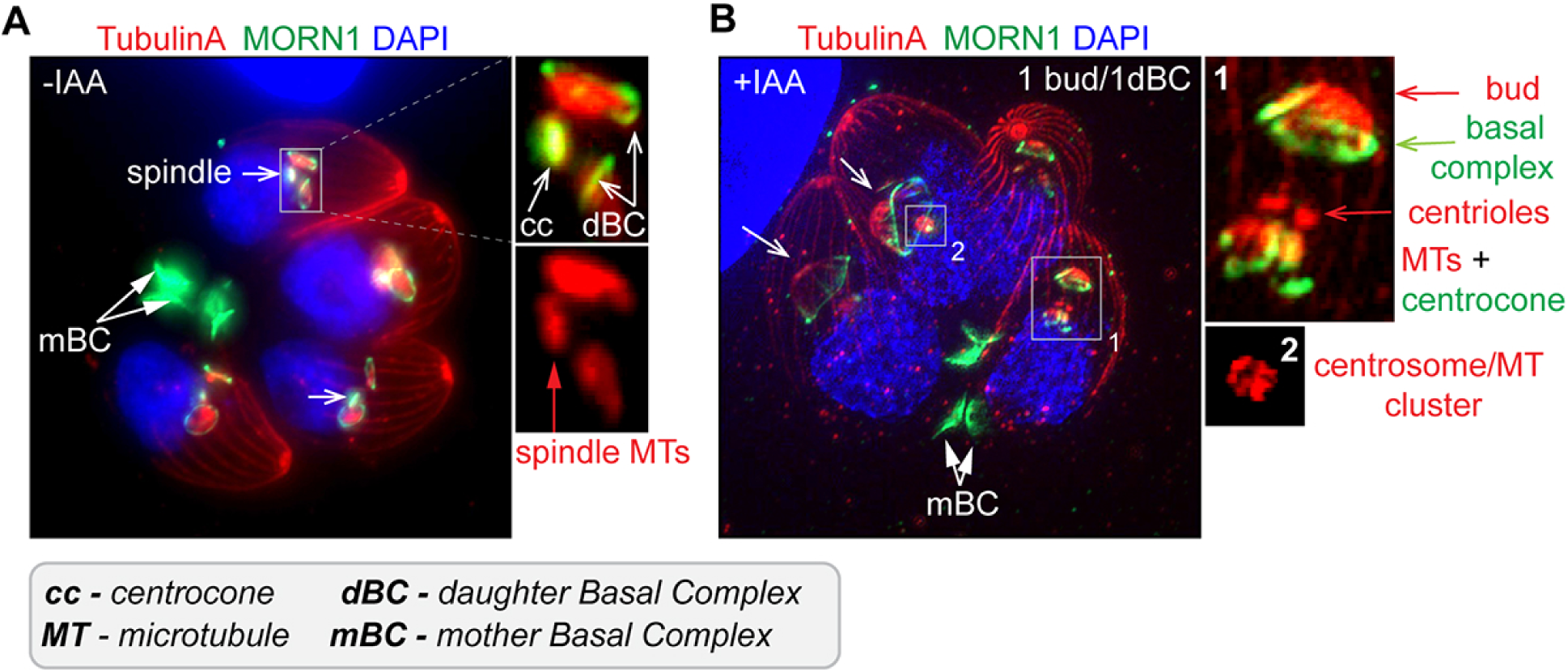
Microscopy analysis of TgCrk4 deficient tachyzoites. (A and B) The ultra-expansion microscopy analysis of RH*ΔKu80TIR1* TgCrk4^AID-HA^ expressing (A) and deficient (B) tachyzoites. Panel A: Co-staining of Tubulin A (α-TubulinA/α-mouse IgG Fluor 568) and TgMORN1 (α-MORN1/α-rabbit IgG Flour 488) shows spindle microtubules located in the extended centrocone and two daughter basal complexes (dBC) encircling subpellicular microtubules of the daughter bud. Nucleus stained with DAPI (blue). Panel B: co-staining of the TgCrk4 deficient parasites (4 h, 500μM IAA) shows assembly of a single daughter bud.

**Figure S4.**
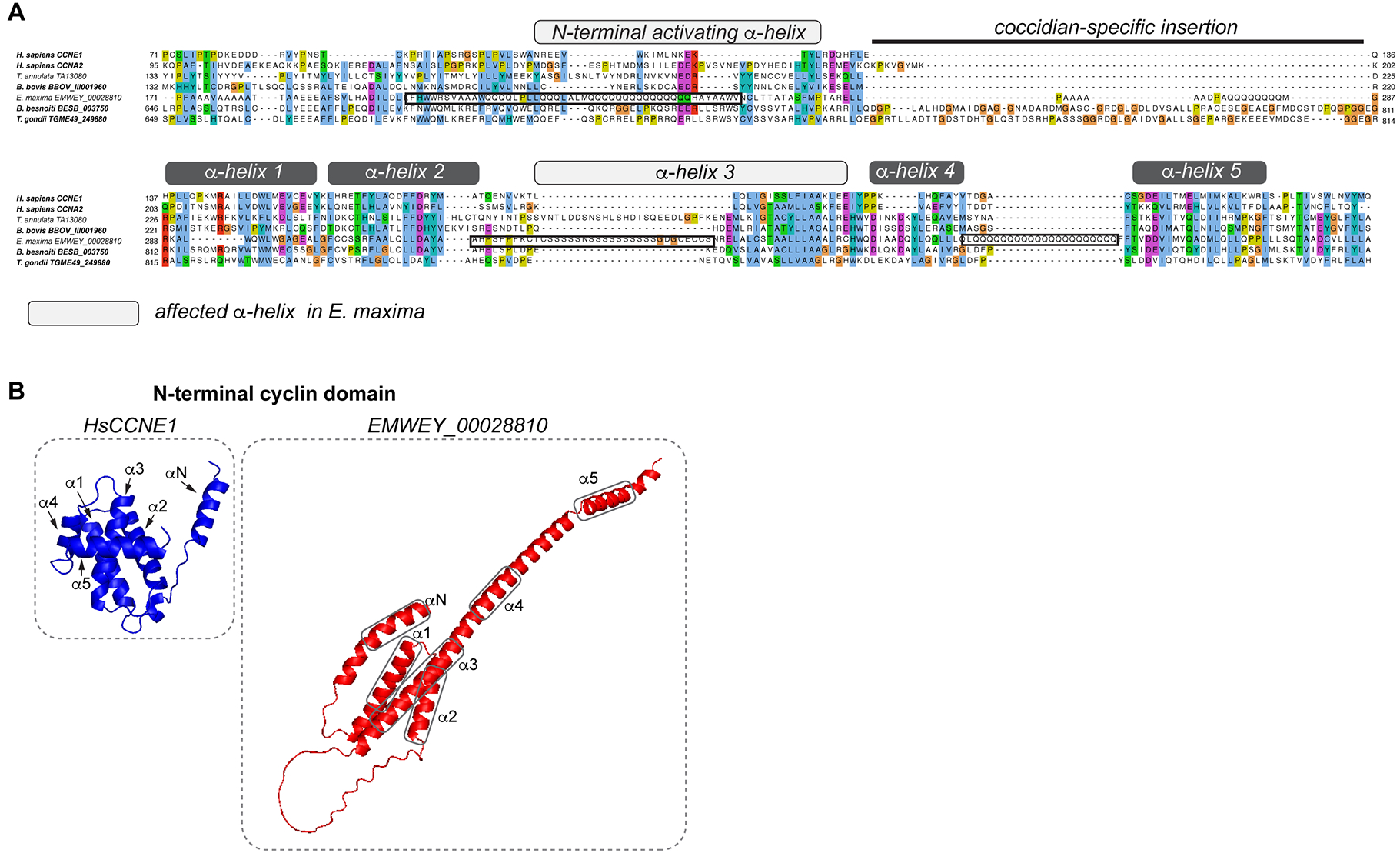
Structural analysis of TgCyc4-related proteins. (A) Alignment of the N-terminal cyclin-domains of apicomplexan Cyc4 proteins with *H. sapiens* Cyclin A2 and Cyclin E1 (MUSCLE). (B) Predicted folding of the cyclin domains of *H. sapiens* Cyclin E1 and Eimeria Cyc4.

**Figure S5.**
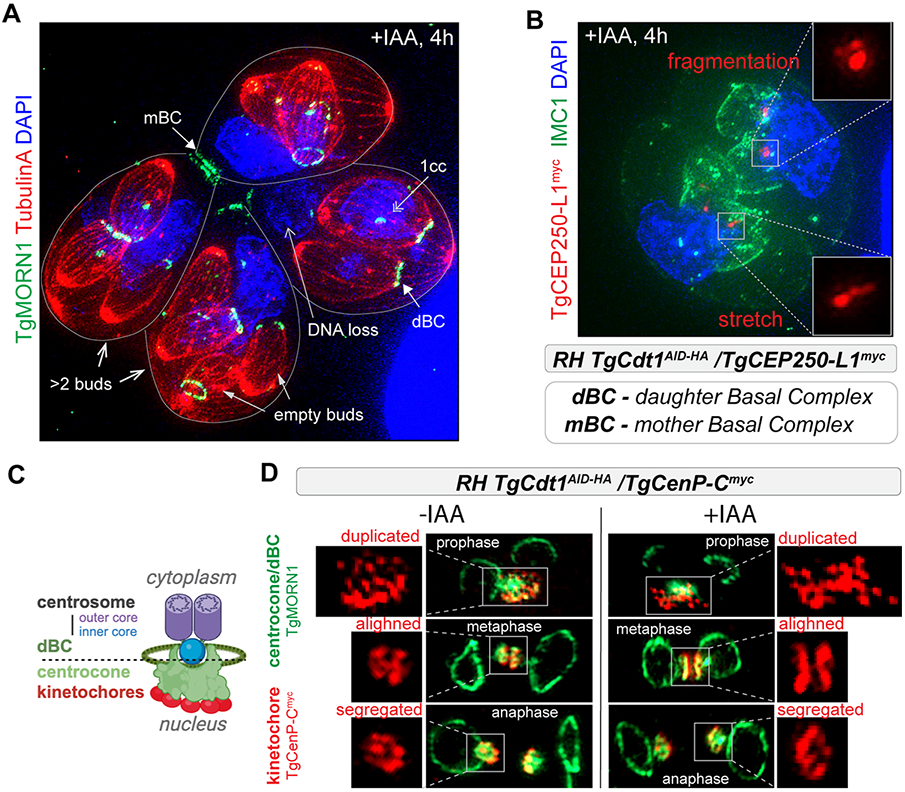
The phenotypic analysis of the TgCdt1 deficient tachyzoites. (A) Ultra-expansion microscopy analysis of RH*ΔKu80TIR1* TgCdt1^AID-HA^ deficient tachyzoites. Staining with Tubulin A (α-TubulinA/α-mouse IgG Fluor 568) visualizes the mother and the daughters’ subpellicular microtubules (buds). The TgMORN1 (α-MORN1/α-rabbit IgG Flour 488) staining shows changes to centrocone and segregates the mother (mBC) and the daughter basal complexes (dBC). DNA mis-segregation defect is depicted with nuclear DAPI stain (blue). (B) The ultra-expansion microscopy images of RH*ΔKu80TIR1* TgCdt1^AID-HA^ deficient tachyzoites expressing TgCEP250-L1^myc^. The inner core of the centrosome was detected with α-myc (α-rabbit IgG Flour 568) antibodies, the parasite surface with α-IMC1 (α-mouse IgG Fluor 488) and nucleus with DAPI stain. The inner core changes caused by TgCdt1 depletion (+IAA, 4 hours) are highlighted in the insets. (C) Schematics of the *T. gondii* perinuclear structures including the bipartite centrosome, centrocone and kinetochores. The drawing depicts one half of the mitotic figure. The dotted line separates nucleoplasm and cytoplasm. dBC – daughter basal complex. (D) The ultra-expansion microscopy images of RH*ΔKu80TIR1* TgCdt1^AID-HA^ tachyzoites expressing TgCenP-C^myc^ after 4 hours incubation without or with IAA. Three stages of mitosis are shown. To determine the relative position of kinetochores and centrocone, the samples were co-stained with α-myc (α-mouse IgG Flour 568) antibodies and α-TgMORN1 (α-rabbit IgG Fluor 488).

